# SPT6 loss Permits the Transdifferentiation of Keratinocytes into an Intestinal Fate that Recapitulates Barrett’s Metaplasia

**DOI:** 10.1101/2021.05.06.442930

**Authors:** Daniella T. Vo, MacKenzie R. Fuller, Courtney Tindle, Mahitha Shree Anandachar, Soumita Das, Debashis Sahoo, Pradipta Ghosh

**Affiliations:** Department of Pediatrics, University of California San Diego; Department of Cellular and Molecular Medicine, University of California San Diego; HUMANOID Center of Research Excellence (CoRE), University of California San Diego; Department of Pathology, University of California San Diego; Department of Computer Science and Engineering, Jacob’s School of Engineering, University of California San Diego; Moore Comprehensive Cancer Center, University of California San Diego; Department of Medicine, University of California San Diego

**Keywords:** Transdifferentiation, Barrett’s metaplasia, keratinocyte, TP63

## Abstract

Transient depletion of the transcription elongation factor SPT6 in the keratinocyte has been recently shown to inhibit epidermal differentiation and stratification; instead, they transdifferentiate into a gut-like lineage. We show here that this phenomenon of *transdifferentiation* recapitulates Barrett’s metaplasia, the only human pathophysiologic condition in which a stratified squamous epithelium that is injured due to chronic acid reflux is trans-committed into an intestinal fate. The evidence we present here not only lend support to the notion that the keratinocytes are the cell of origin of Barrett’s metaplasia, but also provide mechanistic insights linking transient acid exposure, downregulation of SPT6, stalled transcription of the master regulator of epidermal fate TP63, loss of epidermal fate and metaplastic progression. Because Barrett’s metaplasia in the esophagus (BE) is a pre-neoplastic condition with no preclinical human models, these findings have a profound impact on the modeling Barrett’s metaplasia-in-a-dish.

**GRAPHIC ABSTRACT:** 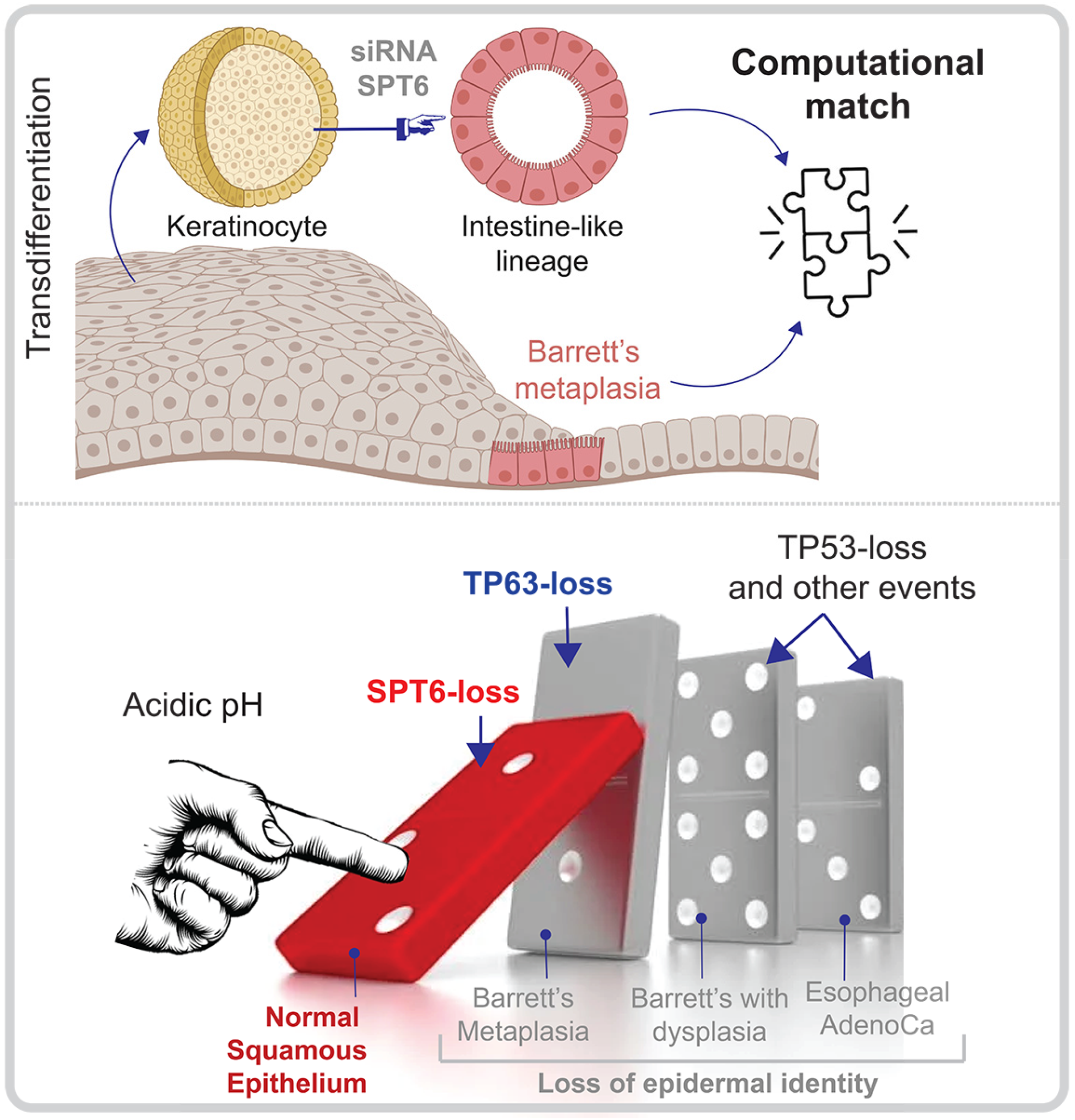

**HIGHLIGHTS:**

- Keratinocytes transdifferentiate into the gut lineage upon depletion of SPT6
- Such transdifferentiation recapitulates Barrett’s metaplasia, not the healthy gut
- Acid downregulates SPT6, which derails the expression and functions of TP63
- Such downregulation precedes the metaplasia-dysplasia-neoplasia cascade

## INTRODUCTION

The stratified squamous epithelium, which is comprised mainly of keratinocytes, acts as a physical barrier and is replaced every few weeks by resident stem cells residing in the basal layer^1,2^. Besides our skin, stratified squamous epithelia form barriers to antigens in the oral cavity and oral pharynx including the palatine and lingual tonsils, the anal canal, the male foreskin, and the female vagina and ectocervix. Recently, it has been shown^3^ that in epidermal stem and progenitor cells, approximately a third of the genes that are induced during differentiation already contain stalled Pol II at the promoters which is then released into productive transcription elongation upon differentiation. Using a combination of Pol II ChIP Seq and RNAi screen, SPT6 was identified as one of the critical mediators of such elongation^3^. SPT6-depleted keratinocytes fail to differentiate into stratified squamous epithelium; instead, they transdifferentiate into a ‘intestine-like’ lineage (by morphology and gene expression analysis; **Figure 1A-B**). This claim of transdifferentiation was supported in part by morphological characteristics in 3D growth and in a more definitive way by transcriptomic studies (<u>GSE153129</u>)^3^. The list of genes that were upregulated ≥ 10-fold in small interfering RNA (siRNA)-depleted SPT6 samples (SPT6i) compared to controls (CTLi). The resultant SPT6-depleted 472-gene signature was used to query the Human Gene Atlas and ARCHS^4^; the latter is a web-based resource that provides access to human and mouse transcriptomic datasets from GEO and SRA^3^. Mechanistically, depletion of SPT6 resulted in stalled transcription of the master regulator of epidermal fate *p63*^4-7^. Studies in SPT6-depleted keratinocytes that were subsequently rescued with exogenous expression of *p63* suggested that SPT6 favors the differentiation into stratified squamous epithelium and arrests the intestinal phenotype through the control of transcriptional elongation of *p63* and its targets. Despite the mechanistic insights into how SPT6 regulates keratinocyte fate, the translational relevance of the observed transdifferentiation of keratinocytes into an intestinal fate remained unknown.

**Figure 1.**
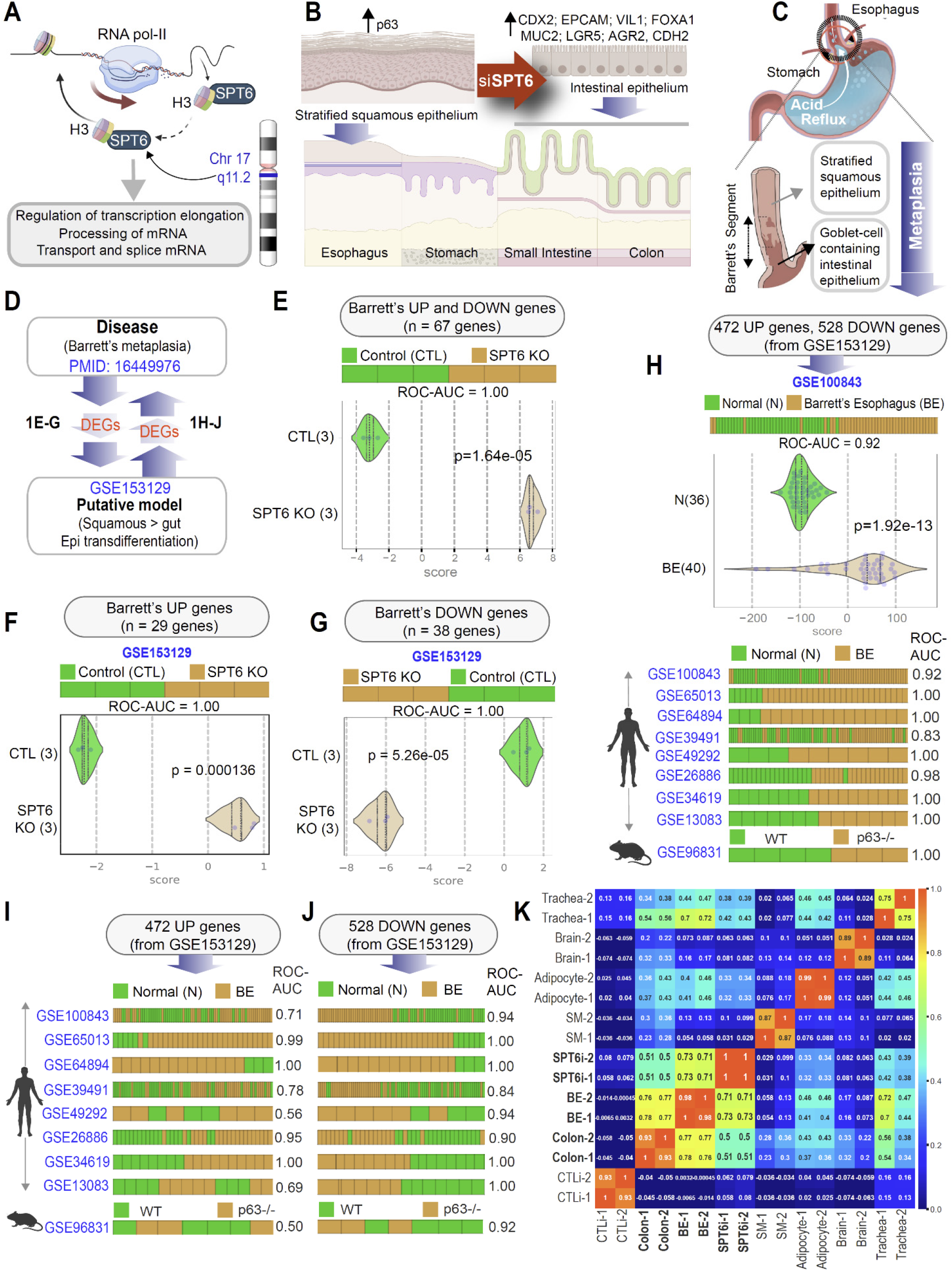
Keratinocyte stem cells depleted of SPT6 trans-differentiates into gut lineage that resembles Barrett’s metaplasia. **A**. Schematic summarizes the chromosomal location of SPT6 and its known functions in transcriptional elongation and mRNA processing. SPT6 coordinates nucleosome dis- and re-assembly, transcriptional elongation, and mRNA processing. SPT6 is a conserved factor that controls transcription and chromatin structure across the genome. **B**. Schematic summarizing the key findings in gene expression and epithelial morphology observed and reported by Li et al.^3^, upon depletion of SPT6 in keratinocyte stem cells by siRNA^3^. While control keratinocytes formed stratified squamous epithelium, siRNA mediated transient depletion of SPT6 in keratinocytes (SPT6i) grew as ‘intestine-like’ monolayers. **C**. Schematic showing the only known human pathophysiologic context in which stratified squamous epithelium is known to be replaced by ‘intestine-like’ epithelium. **D**. Summary of computational approach used in panels 1E-J. **E-G**. Differentially expressed genes (DEGs; see **Supplementary Table 1**) in Barrett’s metaplasia vs. normal esophagus were used to rank order control (CTLi) and SPT6-depleted samples (SPT6i), either using UP genes alone (F), DOWN-genes alone (G) or both UP and DOWN signatures together (E). Results are presented as bar plots. ROC-AUC in all cases reflects a perfect strength of classification (1.00). **H-J**. Differentially expressed genes (DEGs; see **Supplementary Table 2**) between control (CTLi) and SPT6-depleted (SPT6i) samples were used to rank order normal (N) from Barrett’s esophageal (BE) samples across 8 publicly available independent cohorts, either using UP genes alone (I), DOWN-genes alone (J) or both UP and DOWN signatures together (H). See also **Figure S1** for Violin plots for each dataset. ROC-AUC in in each case is annotated on the right side of the corresponding bar plots. **K**. Pearson correlation matrix showing clustering of control (CTLi) and SPT6-depleted (SPT6i) gene expression signatures with brain, colon, BE, adipocyte, trachea, and skeletal muscle. Two distinct RNA-Seq samples of adult origin are shown for each tissue.

Among the various organs that are protected by stratified squamous epithelium, the only human pathophysiologic condition in which a stratified squamous epithelium in adults transdifferentiates into intestinal fate is that described in the foregut, a phenomenon termed Barrett’s metaplasia^8^ of the esophagus (BE). BE develops when the non-keratinized stratified squamous epithelium in the lower esophagus is replaced by a single layer of ‘intestine-like’ cells after a prolonged phase of injury due to chronic acid reflux. The origin of BE remains widely debated; theories include a direct origin from the esophageal stratified squamous epithelium, or by proximal migration and subsequent intestinalization of the gastric epithelium^8-12^. Alternative proposals include a niche cell at the squamocolumnar junction, or cells lining the esophageal gland ducts, or circulating bone-marrow-derived cells^10^. Much of these theories originate from experimental models, and to date, there are no models that recapitulate the process of transdifferentiation of the epithelial lining that is the pathognomonic feature of BE. In fact, our inability to observe the process of metaplastic conversion *in vivo* and the lack of reliable physiological models^13^ are cited as the factors limiting our ability to trace the cell of origin for BE. Despite the lack of models, or dispute surrounding the origin of BE, what is undisputed is that it represents a *bona fide* preneoplastic state; patients with BE have approximately 40-125 times higher risk of esophageal adenocarcinoma than the general population^14^. We hypothesized that the phenomenon of transdifferentiation from stratified squamous to an intestinal fate in SPT6-depleted keratinocytes may resemble and recapitulate the fundamental molecular and cellular aspects of keratinocyte transcommitment in BE.

## RESULTS

### SPT6 loss resembles Barrett’s esophagus

We carried out a comprehensive bidirectional analysis: differentially expressed genes (DEGs), both, up- and downregulated genes^15^ in BE *vs*. normal esophagus (**Figure 1C**) were used to rank order the control *vs*. SPT6-depleted samples, and conversely, DEGs in control *vs*. SPT6-depleted samples were analyzed in all BE datasets publicly available on NCBI as of February 1, 2021 (**Figure 1D**). We found that the combined DEGs (up- and downregulated genes in BE^15^; **Figure 1E; Supplementary Table 1**) as well as the individual up- and downregulated genes (**Figure 1F-G**) were able to independently classify the control and SPT6-depleted samples with perfection (ROC AUC 1.00). The converse was also true, i.e., the combined DEGs (up- and downregulated genes in SPT6-depleted samples; **Figure 1H, Supplementary Table 2**) as well as the individual up- /downregulated signatures (**Figure 1I-J**) were independently able to classify the normal esophageal and BE samples across several independent human datasets. Downregulated genes consistently performed better (ROC AUC ranges from 0.56 – 1.00 in UP-genes, I, and 0.84 - 1.00 in DOWN-genes, J). The DEGs from the SPT6-depleted samples also perfectly classified the BE samples derived from mice lacking *p63* (**Figure 1H-J**); *p63*-/- mice are the only genetic model of BE known to date^16^. Finally, a Pearson correlation matrix revealed that SPT6-depleted samples clustered much closer to BE tissues than to colon (correlation coefficient 0.71-0.73 to BE *vs*. 0.5-0.51 to colon; **Figure 1K**). These findings demonstrate that the transcriptional profile of the SPT6-depleted keratinocyte is more like BE than colon or any other tissue type tested.

### SPT6 loss resembles intestinal metaplasia, not healthy intestinal differentiation

Prior studies have linked loss of *p63*, the master regulator of keratinocyte proliferation and differentiation into a stratified lining^4-7^, as a state that is permissive to the transdifferentiation of stratified squamous epithelium into ‘intestine-like’ metaplasia. In *p63-/-* mice, the stratified lining of both trachea and esophagus are replaced by a highly ordered, columnar ciliated epithelium that is deficient in basal cells^17^. In the same mice, under conditions of programmed damage to the esophageal lining, progenitor cells at the gastroesophageal junction serve as precursors of Barrett’s metaplasia^16^. SPT6 loss in keratinocytes was also associated with a functional loss of *p63*^3^; without SPT6, levels of *p63* protein were diminished, and *p63*-binding sites on the genome were closed, as determined using ATAC seq^3^. Thus, both the SPT6-depleted primary human keratinocyte model and the *p63*-/- mouse model relies upon a final common pathway that escapes an epidermal fate; both lack functional *p63*. We noted that in the *p63-/-* mouse model of BE, Wang et. al, had further delineated that despite the overall similarities, BE segment and intestine tissues have key differences: a set of metaplasia-specific genes is enriched in BE, whereas a set of intestine-specific genes is enriched in the intestine (**Figure 2A; Supplementary Table 4**). Gene set enrichment analyses (GSEA-preranked^18,19^) found these differences also in human BE *vs*. small intestine tissues (GSE13083^20^; **Figure 2B**) and in the SPT6-depleted keratinocyte organoid model reported by Li et al^3^. *vs*. small intestine-derived organoids (**Figure 2C**). Visualization of the same analyses as heatmaps confirm that similar sets of genes within the metaplasia-specific signature was induced in both BE tissue (**Figure 2D**) and in SPT6-depleted (SPT6i) organoids (**Figure 2E**) compared to their respective small intestine-derived samples. These findings further support our argument that SPT6 depletion does not merely trigger intestinal transdifferentiation; it induces metaplasia-specific genes in the setting of de-enrichment of intestine-specific genes.

**Figure 2.**
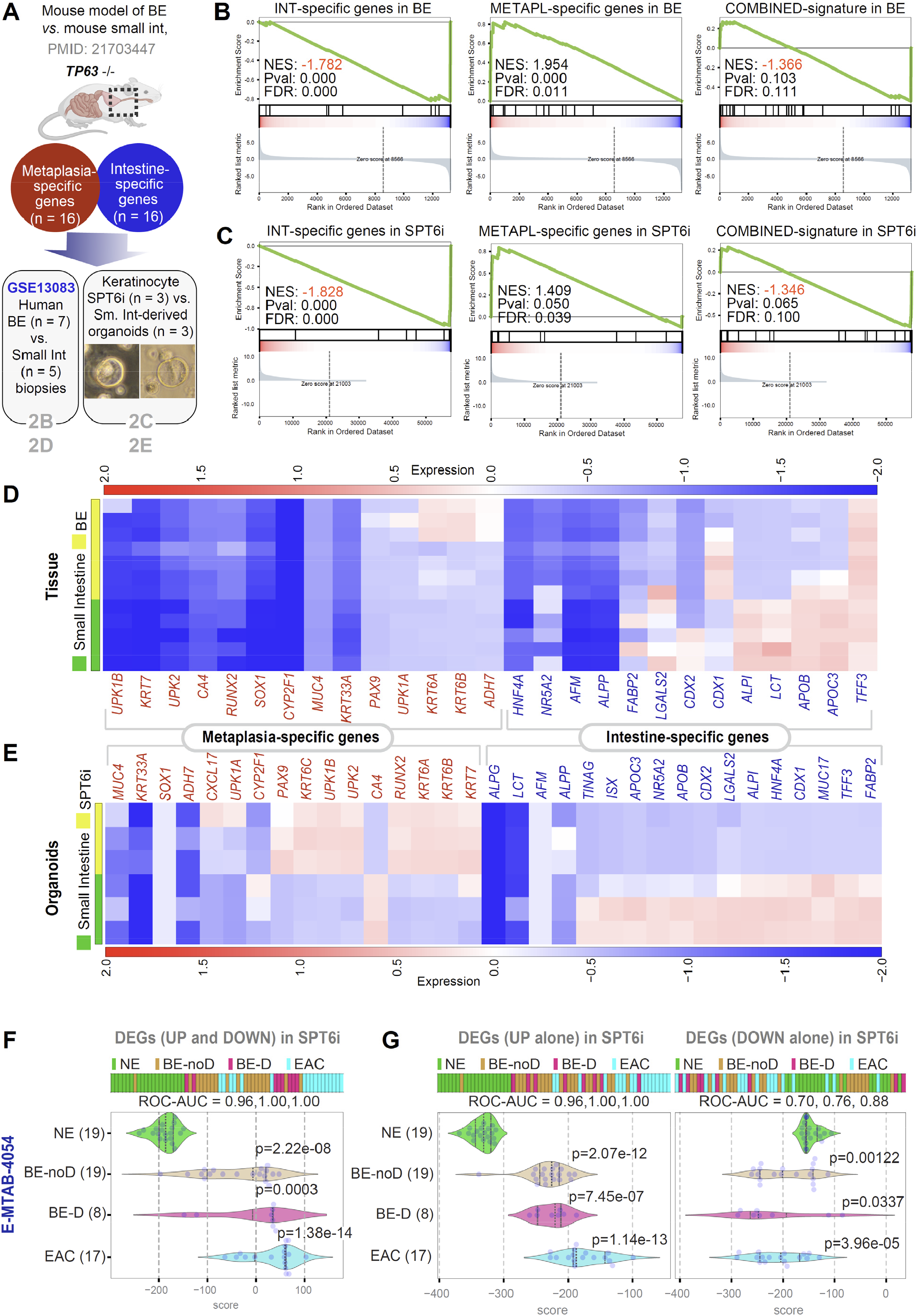
Downregulation of SPT6 enriches metaplasia-specific genes. **A-E**. Schematic in A summarizing the workflow for distinguishing BE from intestine. Using TP63-/- as a strategy to induce BE in mice, a prior study showed that compared to intestinal tissues, BE tissue was enriched in a 16-gene metaplasia-specific signature and de-enriched in a 16-gene intestine-specific signature (see **Supplementary Table 4** for the list of genes). These gene sets were analyzed for enrichment (Gene Set Enrichment Analysis – GSEA pre-ranked analysis) in human BE vs. small intestine tissues (<u>GSE13083</u>; B) and SPT6-depleted (SPT6i) vs. small intestine-derived organoids (C). Heatmaps (D-E) display the levels of expression of the individual metaplasia-specific and intestine-specific genes in the BE vs. small intestine tissues (D) and organoids SPT6-depleted keratinocyte (SPT6i) vs. small intestine-derived organoids (E). **F-G**. Differentially expressed genes (DEGs; see **Supplementary Table 2**) between control (CTLi) and SPT6-depleted (SPT6i) samples were used to rank order normal squamous esophagus (NE) from non-dysplastic Barrett’s esophagus (BE-noD), dysplastic BE (BE-D) and esophageal adenocarcinoma (EAC; n = 12) samples in a RNA seq dataset [<u>E-MTAB-4054</u>]^21^, either using UP genes alone (G; left), DOWN-genes alone (G; right) or both UP and DOWN signatures together (F). Numbers in parenthesis indicate the number of samples. ROC-AUC in each case is annotated below the bar plots. Welch’s two sample unpaired t-test is performed on the composite gene signature score to compute the p values. In multi-group setting each group is compared to the NE control group and only significant p values are displayed.

We next asked if the metaplastic BE-like signature that is induced upon SPT6 depletion stays induced during the progression of BE to esophageal adenocarcinoma (EAC). To this end, we analyzed the DEGs in SPT6-depleted keratinocytes (SPT6i) in a RNA seq dataset comprised of 51 tissue samples, which included normal squamous esophagus, BE metaplasia without dysplasia, BE with low-grade dysplasia (LGD) and esophageal adenocarcinoma (EAC) [<u>E-MTAB-4054</u>]^21^; these were collected at endoscopy from 44 patients. We found that the combined DEGs (up- and downregulated genes) in SPT6-depleted samples classified normal esophagus nearly perfectly (ROC AUC 0.96) from BE samples and perfectly (ROC AUC 1.00) from both dysplastic BE and EAC samples (**Figure 3F**). When we analyzed the upregulated genes (**Figure 3G**; *left*) or the downregulated genes (**Figure 3G**; *right*) separately, we found that the individual up-/downregulated signatures were independently able to classify the normal esophageal samples from all other esophageal tissues representative of progression through the metaplasia-dysplasia-neoplasia cascade. These findings suggest that the BE-like gene expression pattern we observe in SPT6-depleted organoids is conserved during subsequent progression of BE to EAC.

**Figure 3.**
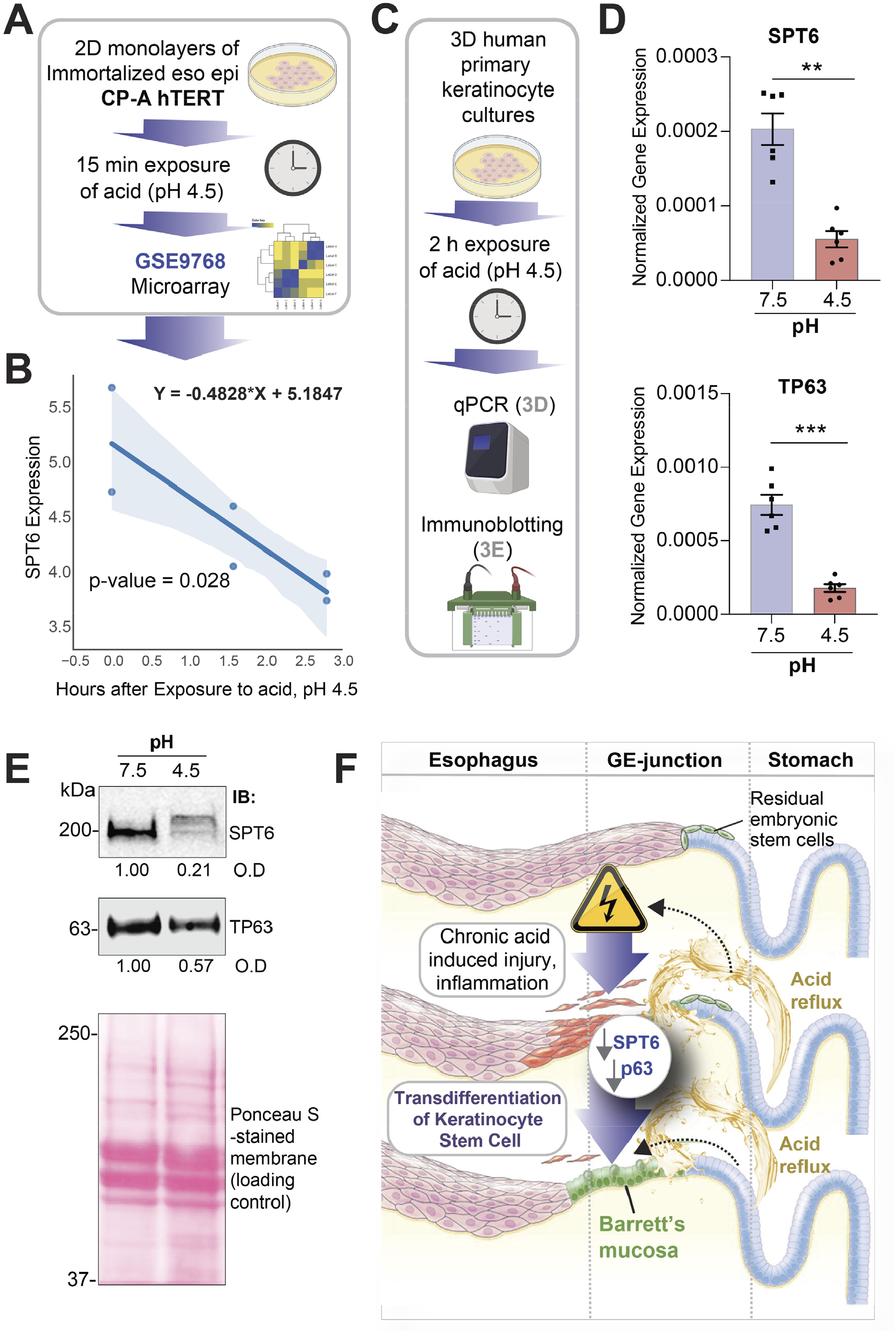
Downregulation of SPT6 can be triggered by exposure to acid. **A**. Publicly available microarray dataset from esophageal epithelial cells (immortalized with hTERT) treated with pH 4.5 for 2 and 6 h were analyzed for SPT6 expression. **B**. Graph displays the abundance of SPT6 transcripts; the x-axis represents the log2(X + 1) transformation of unexposed (0), 2 and 6 hours after exposure to pH 4.5 and the y-axis represents log2 normalized SPT6 expression. Significance was determined by linear regression, where Y = -0.4828*X + 5.1847 and p = 0.028. **C-E**. Schematic in C shows the workflow for the analyses we did here on human primary keratinocytes in 3D culture, after transiently exposing them to acid. Bar graphs in D display the relative expression of SPT6 (top) and TP63 (bottom) in primary keratinocytes exposed to pH 7.5 or 4.5, as determined by qPCR. Error bars represent S.E.M. Significance as determined by t-test, p = ** <0.01; *** < 0.001. n = 3. Immunoblots (IB) in E display the abundance of SPT6 and TP63 proteins in equal aliquots (50 µg) of whole cell lysates of the keratinocytes. O.D = optical density, as determined by band densitometry. Representative immunoblots from 3 independent repeats is shown. **G**. Schematic summarizing the evidence we present here, showing the keratinocyte stem cell as the cell of origin of BE; upon chronic acid (low pH) injury, SPT6 is downregulated in the keratinocyte stem cells. SPT6 suppression, either due to acid exposure (physiologic) or transiently with siRNA (experimentally), causes the spontaneous transdifferentiation of epidermal cells into Barrett’s metaplasia. Li et al.^3^, demonstrated that such transdifferentiation was due to the stalled transcription of the master regulator of epidermal fate p63.

### Acid exposure suppresses SPT6 in keratinocytes

Because BE is a consequence of prolonged acid exposure^8^, we next asked what, if any, might be the impact of low pH on the levels of SPT6 expression in keratinocytes. We found that SPT6 transcripts are downregulated in immortalized esophageal keratinocytes exposed to acid (**Figure 3A-B**). Prior work has also shown that exposure of esophageal keratinocytes to bile and acid causes a reduction in *p63*^22^. When we exposed primary keratinocytes (same cell line used by Li et al.^3^) to pH 4.5 (**Figure 3C**), we found that both SPT6 and TP63 transcripts were reduced (**Figure 3D**). Most importantly, reduced SPT6 and TP63 transcripts in these cells translated to reduced SPT6, and to a lesser extent, TP63 proteins upon acid challenge (**Figure 3E**). These findings help link the SPT6→p63 mechanism(s) outlined by Li et al.^3^, to one of the most definitive physiologic triggers of BE, i.e., exposure to acid.

## Conclusion

Our findings augment the impact of the discoveries reported by Li et al.^3^, in three ways: (i) *First*, they validate transient SPT6-depletion in keratinocytes as an effective strategy for studying origin of BE. Although organoids derived from segments of established BE have been successfully grown in long-term cultures^23^, attempts to model the initiation of BE had thus far been unsuccessful^24^. (ii) *Second*, they weigh in on the long-standing debate surrounding the cell of origin in BE. Controversies exist as to whether BE results from a direct conversion of differentiated cells *via* a process called transdifferentiation, or whether BE develops from niche stem or progenitor cells at the gastroesophageal junction^25,26^. The evidence presented by Li et al.^3^ argues strongly for keratinocyte transdifferentiation or transcommitment as the mechanism. (iii) *Third*, our findings, together with the evidence provided by Li et al.^3^ suggest a mechanism for initiation of BE while being able to connect the dots between physiological triggers of the disease, i.e., chronic acid reflux (**Figure 3F**). For example, acid exposure has been shown to promote intestinal differentiation in both BE explants^27^ and BE-derived adenocarcinoma cell lines^28^, and here we show that acid exposure reduces SPT6 mRNA and protein. Another intriguing coincidence is that SPT6 is located on the long arm of Chr 17q11.2, and loss of heterozygosity (LOH) at this locus is frequently encountered in BE-associated adenocarcinomas^29,30^, representing one of the most frequent LOH in BE^31^. Notably, loss of p53, which is located on the short arm of Chr 17p13, signals risk for BE to adenocarcinoma progression^32^. Unlike the timing of loss of tp53, which happens later in BE-to-cancer progression, microdeletions in the distal long arm of Chr 17q has been proposed as early event in BE initiation^33^. Because microsatellites might be sensitive indicators of disrupted mechanisms and indicate a propensity to mutagenesis, we speculate that microdeletions at Chr17q11.2 could impact SPT6 expression and trigger the transdifferentiation of keratinocytes into BE. It is possible that such microdeletions are a consequence of DNA damage due to repetitive acid injury.

While the strength of our study lies in the rigorous computational validation of one disease model (SPT6-depleted keratinocyte) against diseased tissues from diverse cohorts, there are some notable limitations. For example, although we showed that the impact of suppressing the SPT6→TP63 axis on the gene expression pattern is widely reflected in most if not all BE datasets, whether SPT6 suppression itself is a major and widely prevalent trigger event in BE initiation remains to be established. Similarly, acid exposure reduced both SPT6 mRNA and protein levels, but head-to-head comparisons of the genetic (SPT6-depletion) and physiologic (repetitive acid challenge) triggers need to be analyzed systematically using gene and pathway overlap assessments to fully understand which model is closest to BE and what role SPT6 depletion may plan in its initiation. Finally, prolonged acid injury was not attempted; such studies in conjunction with genomic and epigenomic studies will be insightful to understand how acid injury may lead to SPT6 loss and/or suppression.

## Methods

Detailed methods for computational and experimental approaches are presented in Supplementary Online Materials.

## Acknowledgements

We thank George Sen for transparent communications and for providing access to reagents. This paper was supported by NIH AI141630 and CA100768 (to P.G), GM138385 (to D.S) and AI155696, UG3TR003355 and UG3TR002968 (to D.S, P.G and S.D).

## Author contributions

D.V carried out the computational analysis, with supervision from D.S and P.G. M.F and C.T carried out the studies on 3D organoids under the supervision of S.D and P.G. D.S., S.D. and P.G. provided transdisciplinary expertise, resources, and wrote and edited the manuscript. P.G conceived and directed the project.

## Data availability

All raw data is provided.

## Code availability

The source code is available at https://github.com/sahoo00/BoNE. A bash script scr-be is provided to download all the datasets from our Hegemon web server using a perl script. A Jupyter notebook BE-Analysis.ipynb is provided to perform the analysis and generate the figures in this manuscript.

