## Supplementary Online Materials for "SPT6 loss Permits the Transdifferentiation of Keratinocytes into an Intestinal Fate that Recapitulates Barrett’s Metaplasia"

#### SUPPLEMENTAL ONLINE MATERIALS

#### INVENTORY OF SUPPLEMENTARY MATERIALS

1. TRANSPARENT METHODS
2. SUPPLEMENTARY TABLES
3. SUPPLEMENTARY FIGURES AND LEGENDS
4. REFERENCES CITED

### TRANSPARENT METHODS

1. **Key Resource Table**
2. **Contact for Reagent and Resource Sharing**
3. **Experimental Model and Subject Details**
    1. Keratinocyte organoids
    2. Human intestinal organoids
4. **Method Details**
    1. Computational methods
        1. Gene expression databases
        2. Boolean analysis
        3. Generation of gene signature codes
        4. Measurement of classification strength or prediction accuracy
        5. Test and validation of Barrett's esophagus datasets
        6. Correlation analysis
        7. GeneSet Enrichment analysis (GSEA)
        8. Statistical Analyses
    2. Experimental methods
        1. Human epidermal keratinocyte culture
        2. Isolation and culture of human small intestine organoids
        3. Exposure to acid injury
        4. RNA isolation
        5. Quantitative (q)RT-PCR
        6. Quantitative immunoblotting
5. **Quantification and Statistical Analysis**
    1. Statistical Analysis
    2. Replications
6. **Data and Software Availability**
    1. The source code can be accessed at https://github.com/sahoo00/BoNE.

#### KEY RESOURCE TABLE:

### MATERIALS & REAGENTS

#### ANTIBODIES (USED FOR IMMUNOBLOTS)

| Name | Manufacturer | Catalog number | Dilution factor |
|---|---|---|---|
| **Anti-SPT6** | Thermo Fisher Scientific | A300-801A | 1:500 |
| **Rabbit polyclonal anti-β-tubulin** | Santa Cruz Biotechnology | sc-9104 | 1:500 |
| **Mouse monoclonal anti-GAPDH** | Santa Cruz Biotechnology | sc-365062 | 1:500 |
| **IRDye 800CW Goat anti-Mouse IgG Secondary** | LI-COR Biosciences | 926-32210 | 1:10,000 |
| **IRDye 680RD Goat anti-Rabbit IgG Secondary** | LI-COR Biosciences | 926-68071 | 1:10,000 |

#### BIOLOGICAL SAMPLES AND CELL LINES

| Model | Source | Citation/Catalog# |
|---|---|---|
| **Human Epidermal Keratinocyte Culture (from neonatal foreskin)** | UC San Diego HUMANOID Center of Research Excellence | Li et al.[1] |
| **Human intestinal organoids** | | Sharma et al.[2] |
| **L-WRN cells** | ATCC | CRL-3276 [3] |

#### PRIMER SEQUENCES

| Target human genes | Forward/Reverse | Primer sequence |
|---|---|---|
| SPT6 | Forward | CCGTGTCCACCCTGAGAC |
| | Reverse | CATAGCCCTGCCTCTCCA |
| TP63 | Forward | GACAGGAAGGCGGATGAAGATAG |
| | Reverse | TGTTTCTGAAGTAAGTGCTGGTGC |
| 18S | Forward | GTAACCCGTTGAACCCCATT |
| | Reverse | CCATCCAATCGGTAGTAGCG |

#### INSTRUMENTS

| | | |
|---|---|---|
| **Countess II Automated Cell Counter** | Thermo Fisher Scientific | AMQAX1000 |
| **Canon Rebel XS DLSR** | Canon | |
| **MiniAmp Plus Thermal Cycler** | Applied Biosystems | A37835 |
| **QuantStudio5** | Applied Biosystems | A28140 |
| **Light Microscope (brightfield images)** | Carl Zeiss LLC | Axio Observer, Inverted; 491917-0001-000 |

#### SOFTWARE

| | |
|---|---|
| **ImageJ** | https://imagej.nih.gov/ij/index.html |
| **GraphPad Prism** | https://www.graphpad.com/scientific-software/prism/ |
| **QuantStudio Design & Analysis Software** | https://www.thermofisher.com/us/en/home/global/forms/life-science/quantstudio-3-5-software.html |

| | | |
|---|---|---|
| **Illustrator (ADOBE)** | https://www.adobe.com/products/illustrator.html | |
| **ImageStudio Lite (LI-COR Sciences)** | https://www.licor.com/bio/image-studio-lite/ | |
| **ENZYMES, CHEMICALS, DISPOSABLES AND REAGENTS** | | |
| **PVDF Transfer Membrane, 0.45µM (for blotting)** | Thermo Scientific | 88518 |
| **PowerUp™ SYBR™ Green Master Mix (for qPCR)** | Applied Biosciences | A25741 |
| **qScript™ cDNA SuperMix (for qPCR)** | QuantaBio | 101414 |
| **Ethanol** | Koptec | UN1170 |
| **Protease inhibitor cocktail (for cell lysis)** | Roche | 11 873 580 001 |
| **Tyr phosphatase inhibitor cocktail (for cell lysis)** | Sigma-Aldrich | P5726 |
| **Ser/Thr phosphatase inhibitor cocktail (for cell lysis)** | Sigma-Aldrich | P0044 |
| **100% Methanol (for priming PVDF membrane)** | Supelco | MX0485 |
| **Glycine** | Fisher Scientific | BP381-5 |
| **Bovine Serum Albumin** | Sigma-Aldrich | A9647-100G |
| **Triton-X 100 (for cell lysis)** | Sigma-Aldrich | X100-500ML |
| **TrypLE Select** | Thermo Scientific | 12563-011 |
| **Advanced DMEM/F-12** | Thermo Scientific | 12634-010 |
| **HEPES Buffer** | Life Technologies | 15630080 |
| **Glutamax** | Thermo Scientific | 35050-061 |
| **Penicillin-Streptomycin** | Thermo Scientific | 15140-122 |
| **Collagenase Type I** | Thermo Scientific | 17100-017 |
| **Matrigel** | Corning | 354234 |
| **B-27** | Thermo Scientific | 17504044 |
| **N-acetyl-L-cysteine** | Sigma-Aldrich | A9165 |
| **Nicotinamide** | Sigma-Aldrich | N0636 |
| **FGF-7 (KGF)** | PeproTech | 100-19-50ug |
| **FGF10** | PeproTech | 100-26-50ug |
| **A-83-01** | Bio-Techne Sales Corp. | 2939/50 |
| **SB202190** | Sigma-Aldrich | S7067-25MG |
| **Y-27632** | R&D Systems | 1254/50 |
| **DPBS** | Thermo Scientific | 14190-144 |
| **Ultrapure Water** | Invitrogen | 10977-015 |
| **EDTA** | Thermo Scientific | AM9260G |
| **Hydrocortisone** | STEMCELL Technologies | 7925 |
| **Heparin** | Sigma Aldrich | H3149 |
| **Fetal Bovine Serum** | Sigma-Aldrich | F2442-500ML |
| **EpiVita Media** | Cell Applications | 141-500a |
| **Animal Component-Free Cell Dissociation Kit** | STEMCELL Technologies | 5426 |
| **Red Blood Cell Lysis Buffer** | Invitrogen | 00-4333-57 |
| **Cell Recovery Solution** | Corning | 354253 |

| | | |
|---|---|---|
| **Sodium Azide (for antibody dilutions)** | Fisher Scientific | S227I-100 |
| **Quick-RNA MicroPrep Kit** | Zymo Research | R1051 |
| **Quick-RNA MiniPrep Kit** | Zymo Research | R1054 |
| **Ethyl alcohol, pure** | Sigma-Aldrich | E7023 |
| **TRI Reagent** | Zymo Research | R2050-1-200 |
| **2x SYBR Green qPCR Master Mix** | Bimake | B21203 |
| **qScript cDNA SuperMix** | Quanta Biosciences | 95048 |
| **Applied Biosystems TaqMan Fast Advanced Master Mix** | Thermo Scientific | 4444557 |
| **18S, Hs99999901_s1** | Thermo Scientific | 4331182 |
| **OTHER** | | |
| **6-well Tissue Culture Plate** | Genesee Scientific | 25-105 |
| **12-well Tissue Culture Plate** | CytoOne | CC7682-7512 |
| **Cell Scraper** | Millipore Sigma | C5981-100EA |
| **Countess Cell Counting Chamber Slides** | Invitrogen | C10312 |
| **Trypan Blue Stain** | Invitrogen | T10282 |
| **70 um Cell Strainer** | Thermo Fisher Scientific | 22-363-548 |
| **100 um Cell Strainer** | Corning | 352360 |
| **RNase Away** | Thermo Fisher Scientific | 14-375-35 |

#### CONTACT FOR REAGENT AND RESOURCE SHARING

Pradipta Ghosh for experimental methods and reagents

Debashis Sahoo for computational methods and datasets

#### DETAILED METHODS:

##### *Computational methods*

**Gene expression databases:** Publicly available microarray and gene expression databases were downloaded from the National Center for Biotechnology Information (NCBI) Gene Expression Omnibus website (GEO) [4-6]. If the dataset is not normalized, RMA (Robust Multichip Average)[7,8] is used for microarrays and CPM (Counts Per Millions)[9,10] (citation) is used for RNASeq data for normalization. We used log2(CPM+1) to compute the final log-reduced expression values for RNASeq data. Accession numbers for these crowdsourced datasets are GSE153129, GSE100843, GSE65013, GSE64894, GSE39491, GSE49292, GSE26886, GSE34619, GSE13083, GSE96831, GSE120795, GSE129153, GSE148818, GSE58963, GSE157059, GSE9768 and GSE70051 and are provided in the figures and manuscript. [11-13].

**Boolean Analysis:** *Boolean logic* is a simple mathematic relationship of two values, i.e., high/low, 1/0, or positive/negative. The Boolean analysis of gene expression data requires first the conversion of expression levels into two possible values. The *StepMiner* algorithm is used to perform Boolean analysis of gene expression data[14]. *Boolean analysis* is a statistical approach that creates binary logical inferences that explain the relationships between phenomena. Boolean analysis is performed to determine the relationship between the expression levels of pairs of genes. The *StepMiner* algorithm is applied to gene expression levels to convert them into Boolean values (high and low). In this algorithm, first the expression values are sorted from low to high and a rising step function is fitted to the series to identify the threshold. Middle of the step is used as the *StepMiner* threshold. This threshold is used to convert gene expression values into Boolean values. A noise margin of 2-fold change is applied around the threshold to determine intermediate values, and these values are ignored during Boolean analysis.

**Generation of gene signature scores:** Gene expression values were normalized according to a modified Z-score approach centered around *StepMiner* threshold (formula = (expr - SThr)/3*stddev). The samples were ordered according to average of the normalized gene expression values in the given gene list. Gene signature score is computed as a linear combination of the normalized gene expression values (the modified Z-score as described above). Samples are ordered using the gene signature score and the strength of the association between gene expression and disease annotation is computed using ROC-AUC measurement. A barplot is used to visualize the sample ordering with different color codes for the sample annotation. Additionally, a set of violin plots is used just below the barplot to demonstrate the distribution of the gene signature score across different sample annotations.

**Measurement of classification strength or prediction accuracy:** Receiver operating characteristic (ROC) curves were computed by simulating a score based on the ordering of samples that illustrates the diagnostic ability of binary classifier system as its discrimination threshold is varied along the sample order. The ROC curves were created by plotting the true positive rate (TPR) against the false positive rate (FPR) at various threshold settings. The area under the curve (often referred to as simply the AUC) is equal to the probability that a classifier will rank a randomly chosen IBD samples higher than a randomly chosen healthy samples. In addition to ROC AUC, other classification metrics such as accuracy ((TP + TN)/N; TP: True Positive; TN: True Negative; N: Total Number), precision (TP/(TP+FP); FP: False Positive), recall (TP/(TP+FN); FN: False Negative) and f1 (2 * (precision * recall)/(precision + recall)) scores were computed. Precision score represents how many selected items are relevant and recall score represents how many relevant items are selected. Fisher exact test is used to examine the significance of the association (contingency) between two different classification systems (one of them can be ground truth as a reference).

**Test and Validation of Barrett's Esophagus Datasets:** A Boolean Network Explorer (BoNE) computational tool was introduced in Sahoo D et al.[15] to model natural progressive time-series changes in major cellular compartments that initiate, propagate and perpetuate inflammation in IBD and are likely to be important for disease progression. BoNE provides an integrated platform for the construction, visualization and querying of a network of progressive changes much like a disease map. A published gene signature for Barrett's Esophagus UP genes and DOWN genes[16] were used to train Boolean models that distinguish SPT6-depleted and Control samples in GSE153129. Another gene signature for SPT6-depleted versus Control (GSE153129) was found using the differentially expressed gene file provided by Li et al.[1] (**Supplementary Table 2**), where genes were considered significant if they had a log fold change of greater than 10 or less than -10 and p-value of less than 0.1. The gene signature for SPT6-depleted versus Control samples (GSE153129) was used train Boolean models that distinguish normal esophagus and Barrett's metaplasia in humans (GSE100843, GSE65013, GSE64894, GSE39491, GSE49292, GSE26886, GSE34619, GSE13083, E-MTAB-4054), or WT (wildtype) versus p63-/- in mice (GSE96831). In this model, a path score is computed as mentioned in section "Generation of gene signature scores" that is used to order the samples. The sample ordering is evaluated using the sample annotation (Normal, Barrett's Esophagus) using ROC-AUC.

**Correlation Analysis:** Correlation analysis was performed using Python pandas.DataFrame.corr (version 1.1.5). Normalized counts (that are not log-reduced) for up- and down- regulated genes (filtered by Fold-change (SPT6i vs. CTLi) > 10) were used to recreate the correlation matrix from Li et al.[1], with the addition of Barrett's metaplasia samples. The analysis includes a selection of samples from the following GEO datasets: GSE153129

(SPT6-depleted and Control), GSE120795 (colon, brain, skeletal muscle), GSE129153 (adipocyte), GSE148818 (trachea) and GSE58963 (BE) Specific sample numbers used to generate the correlation matrix are provided in **Supplementary Table 1**.

**GeneSet Enrichment Analysis (GSEA):** GeneSet Enrichment Analysis (GSEA) was performed using Python gseapy (0.10.2 package). Difference in average expression values of two groups is used to compute gene rank file. Metaplasia-specific and intestine-specific mouse genes from Wang et. al[17] were converted to human genes using BoNE. Intestine-specific genes only, metaplasia-specific genes only, and combined metaplasia-specific and intestine-specific genes correspond to the three genesets tested for Barrett's metaplasia versus small intestine samples (GSE13083), and SPT6-depleted (GSE153129) versus small intestine samples (GSE157059). GSEA pre-ranked analysis is performed on the precomputed rank file to check the significance of geneset enrichment score and generate the enrichment plot. GSEA computes four key statistics for the gene set enrichment analysis report: Enrichment Score (ES), Normalized Enrichment Score (NES), False Discovery Rate (FDR), Nominal P Value.

##### *Experimental methods*

**Human Epidermal Keratinocyte Culture:** SPT6 knock-down and wild-type human epidermal keratinocytes were cultured under 3D spheroid conditions as previously described[18] for colon-derived organoids. Briefly, primary human epidermal keratinocytes derived from human neonatal foreskin were used for all cell culture studies. Cells were seeded in Matrigel (Corning, 354234) domes at 5E4 cells per well in a 24 well plate. To allow for complete polymerization of the Matrigel, the plate was inverted and incubated at 37°C for 10 minutes before proliferation media (50% conditioned media prepared from L-WRN cells (ATCC, CRL-3276[3]) containing Wnt3a, R-spondin, and noggin) or differentiation media (5% conditioned media) was added to each well. Cells were maintained at 37°C/5% CO2 humidified conditions and media was changed every 2-3 days until spheroids were fully formed, and crypt budding was visible.

**Isolation and culture of human small intestine organoids:** Intestinal organoids were isolated and cultured following methods previously outlined by Sharma et al.[2] Briefly, human ileal tissue specimens were digested in collagenase type I at 37°C. Vigorous pipetting was performed every 10 minutes until tissue fragments dissociated into single epithelial units. Subsequently, wash media (DMEM/F12, 1X glutamax, 10% FBS) was added to neutralize the collagenase digestion, and the cell suspension was passed through a 70 µm filter. The cells were seeded in Matrigel domes and cultured in proliferation media in a 37°C/5% CO2 humidified incubator. Media changes were performed every 2 days until organoids formed and reached confluency.

**Exposure to acid injury:** Human epidermal keratinocytes were cultured under 2-D and 3-D conditions in Human EpiVita Media (Cell Applications, 141-500a). Keratinocytes were seeded in monolayers at 2E4 cells per well and in Matrigel at 1E5 cells per well in a 12 well plate. Media changes were performed every 2-3 days. Cultures were maintained at 37°C/5% CO2 humidified conditions for 5-6 days until the monolayers reached confluency, and spheroids formed in the 3D culture. Subsequently, cells were cultured in acidic media with a pH range of 4.5-7.5. The median was prepared by adding 1N HCl drop-wise to EpiVita media until the desired pH was achieved. After 2 hours, the acidic media was removed from the keratinocytes and replaced with fresh EpiVita media. Cells were cultured for an additional 6 hours.

**RNA Isolation:** The Matrigel domes were scraped from the surface of the 12 well plate, and spheroids were collected in cell recovery solution (Corning, 354253) and incubated for 1 h at 4°C under constant rotation.

Monolayers and spheroids were lysed in 200 µl of RNA lysis buffer per well, and RNA Isolation was performed following instructions from the Zymo Research Quick-RNA MicroPrep Kit (R1051).

**Quantitative (q)RT-PCR:** Gene expression in spheroids and monolayers was measured by qRT-PCR using 2x SYBR Green qPCR Master Mix (Bimake, B21203). cDNA was amplified with gene-specific primer/probe set for SPT6 and Barrett's Esophagus markers and qScript cDNA SuperMix 5x (Quanta Biosciences, 95048). qRT-PCR was performed with the Applied Biosystems QuantStudio 5 Real-Time PCR System. Cycling parameters were as follows: 95°C for 20 s, followed by 40 cycles of 1 s at 95°C and 20 s at 60°C. All samples were assayed in triplicate and eukaryotic 18S ribosomal RNA was used as a reference. Primer sequences are provided in Table of reagents (above).

**Quantitative Immunoblotting:** For immunoblotting, keratinocytes protein samples were boiled in Laemmli sample buffer, separated by SDS-PAGE and transferred onto 0.4mm PVDF membrane (Millipore) prior to blotting. Post transfer, membranes were blocked using 5% Non-fat milk or 5% BSA dissolved in PBS. Primary antibodies (anti-SPT6; Thermo Fisher, A300-801A; dilution 1:500) were prepared in blocking buffer containing 0.1% Tween-20 and incubated with blots, rocking overnight at 4°C. After incubation, blots were incubated with secondary antibodies for one hour at room temperature, washed, and imaged using a dual-color Li-Cor Odyssey imaging system.

#### QUANTIFICATION AND STATISTICAL ANALYSIS

**Statistical analyses in computational studies:** All statistical tests were performed using R version 3.2.3 (2015-12-10). Standard t-tests were performed using Python scipy.stats.ttest_ind package (version 0.19.0) with Welch's Two Sample t-test (unpaired, unequal variance (equal_var=False), and unequal sample size) parameters. Linear regression was performed using Python scipy.stats.linregress package (version 1.7.0). Multiple hypothesis correction were performed by adjusting *p* values with statsmodels.stats.multitest.multipletests (fdr_bh: Benjamini/Hochberg principles).[19] Violin, Swarm and Bubble plots are created using Python seaborn package version 0.10.1.

**Statistical analyses in experimental studies and replications:** All experiments were repeated at least three times, and results were presented either as one representative experiment or as average ± S.E.M. Statistical significance was assessed with unpaired Student's t test. For all tests, a p-value of 0.05 was used as the cutoff to determine significance [*p < 0.05, **p < 0.01, ***p < 0.001, ****p < 0.0001]. p-values are indicated in each figure. All statistical analysis was performed using GraphPad prism 8.

#### DATA AND SOFTWARE AVAILABILITY

**Data availability:** All raw data is available for sharing.

**Software and Code availability:** The source code is available at https://github.com/sahoo00/BoNE. A bash script scr-be is provided to download all the datasets from our Hegemon web server using a perl script. A Jupyter notebook BE-Analysis.ipynb is provided to perform the analysis and generate the figures in this manuscript. Software programs (listed in Key Resource Table, above) are all publicly accessible through valid licenses.

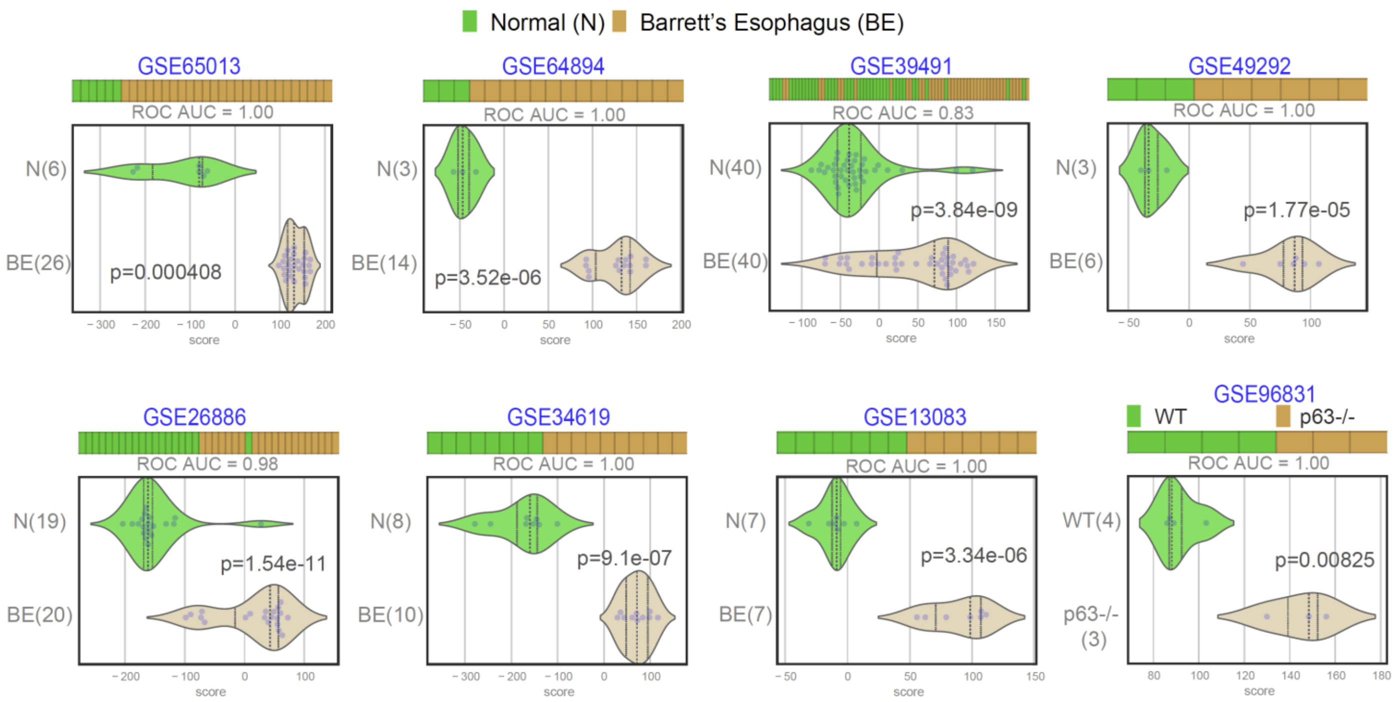

**Supplementary Figure 1.** [Related to **Figure 1**]

**Differentially expressed genes (DEGs) in SPT6-depleted samples recapitulate the altered gene expression patterns seen in multiple independent cohorts of Barrett's esophagus**. DEGs from control vs SPT6-depleted samples in Li et al [1]. were used to rank order normal (N) from Barrett's esophageal (BE) samples across 8 publicly available independent cohorts. ROC AUC in in each case is annotated.

### SUPPLEMENTARY TABLES

**Supplementary Table 1** [Related to **Figure 1D-G**]

**Barrett's Metaplasia Gene Signature** **[PMID: 16449976]**

**Ref:** https://www.nature.com/articles/1209357/tables/1

| **UP Genes** | | **DOWN Genes** | | |
|---|---|---|---|---|
| TNFRSF10C | AZGP1 | MAL | FDXR | NEDD9 |
| LTA | CREB3L1 | LGALS7 | CDA | ANXA8 |
| ARPC3 | FOXA3 | RELN | GPX3 | SERPINB3 |
| GAL5 | TCEAL1 | ANXA1 | CBR3 | SPRR2C |
| TSPAN-1 | NR0B2 | ARS | SULT2B1 | CSTA |
| LAMC2 | CXCL3 | EMP1 | PAX9 | LY6G6C |
| TS4SF8 | GDF15 | VAT1 | RARG | S100A2 |
| CRIP1 | PDGFA | ALOX12 | TRIM29 | TPD52L2 |
| TM4SF3 | GJB1 | KRT6A | CRABP2 | CSTB |
| KRT8 | INSR | ST3GAL4 | MAFG | KIAA0657 |
| COL4A2 | PNPLA2 | KRT4 | NFRKB | PPL |
| SERPINH1 | AGR2 | PGD | ECM1 | |
| LYZ | AADAT | KRT1 | INI1A | |
| CYBA | | FDXR | ARF4L | |
| HGD | | CDA | | |
| PSMB8 | | GPX3 | | |

**Supplementary Table S1 [**Related to **Figure 1D-G]:** Differentially expressed genes (DEGs) from Barrett's esophagus (BE) *vs*. normal esophageal mucosa (NE) from Wang et al.[16] used to rank order control and SPT6-depleted samples from Li et al[1]. in **Figure 1D-G**.  The p-value was <0.001 between BE vs. NE in analysis of variance (ANOVA). The FDR was <0.001 in significance analysis of microarray (SAM) with more than two-fold changes.

| **Supplementary Table 2** [Related to **Figure 1H-J]** | | | | | | |
|---|---|---|---|---|---|---|
| **SPT6i vs CTLi Genes (Fold-change > 10 or < -10, p-value < 0.1)** | | | | | | |
| **Source:** GSE153129-Supplementary-4.xlsx | | | | | | |
| **UP** | | | | | **DOWN** | |
| **Gene Symbol** | **Fold-change (SPT6i vs. CTLi) > 10** | **p_value** | | **Gene Symbol** | **Fold-change (SPT6i vs. CTLi) < -10** | **p_value** |
| **APOBEC3B-AS1** | 2.13E+07 | 7.03E-06 | | **SNORD140** | -38871200 | 1.03E-07 |
| **CGB5** | 19925900 | 9.97E-05 | | **UGT1A9** | -17082600 | 7.31E-06 |
| **MDC1-AS1** | 1.93E+07 | 2.62E-06 | | **SLAMF8** | -2426260 | 2.34E-08 |
| **FKBP6** | 16680200 | 7.32E-07 | | **RNU2-1** | -1228450 | 0.004197 |
| **TBC1D3P2** | 16114900 | 1.07E-06 | | **MSMP** | -148886 | 0.002621 |
| **CGB3** | 8.01E+06 | 3.26E-07 | | **LOR** | -8425.69 | 6.44E-05 |
| **CYP3A7-CYP3A51P** | 7.74E+06 | 0.000467 | | **CRNN** | -3873.47 | 2.27E-06 |
| **PSG3** | 5164160 | 4.52E-06 | | **LCE3C** | -2843.91 | 1.81E-06 |
| **KCNK4-TEX40** | 1399460 | 0.000588 | | **MUC21** | -2167.64 | 7.26E-06 |
| **ANKRD20A8P** | 1.20E+06 | 5.36E-06 | | **LCE1D** | -1722.82 | 0.000226 |
| **GOLGA8S** | 492416 | 0.000199 | | **LCE1C** | -1702.44 | 6.62E-05 |
| **FOXD4L6** | 238565 | 0.000993 | | **LCE2D** | -1676.54 | 4.95E-05 |
| **P2RY11** | 2.59E+04 | 0.000589 | | **C1orf68** | -1652.5 | 1.24E-05 |
| **GPR37L1** | 4.83E+02 | 0.00172 | | **LCE2B** | -1588.73 | 0.000268 |
| **TNFSF15** | 419.423 | 2.46E-05 | | **KRT1** | -1476.25 | 1.08E-05 |
| **CDHR2** | 276.283 | 3.75E-05 | | **SPRR2G** | -1261.91 | 3.53E-07 |
| **LINC01132** | 2.55E+02 | 0.000561 | | **LCE6A** | -1083.99 | 4.31E-05 |
| **APOBEC3H** | 2.36E+02 | 0.001296 | | **LCE2A** | -1022.03 | 0.000118 |
| **CLDN9** | 2.20E+02 | 1.77E-06 | | **ALOX12B** | -995.863 | 1.60E-05 |
| **APOLD1** | 203.536 | 4.94E-07 | | **CASP14** | -988.923 | 6.99E-06 |
| **TNFRSF10C** | 185.85 | 4.40E-06 | | **PSG7** | -966.6 | 2.28E-05 |
| **LINC00589** | 1.54E+02 | 9.41E-07 | | **LIPM** | -945.041 | 1.38E-05 |
| **FOXA1** | 1.51E+02 | 1.90E-08 | | **CYP4F22** | -814.896 | 6.90E-05 |
| **PRR15** | 1.33E+02 | 0.000228 | | **AADACL2** | -806.028 | 3.60E-05 |
| **DCST2** | 119.559 | 8.59E-06 | | **FLG2** | -791.798 | 9.97E-05 |
| **AGR2** | 117.539 | 9.76E-05 | | **HAL** | -669.462 | 3.90E-06 |
| **ZSCAN4** | 9.58E+01 | 3.70E-05 | | **KPRP** | -651.761 | 0.000131 |
| **CXCL5** | 9.51E+01 | 1.66E-06 | | **LCE1F** | -601.582 | 7.17E-05 |
| **LDB3** | 9.07E+01 | 8.01E-05 | | **PLA2G3** | -568.464 | 4.57E-06 |
| **SLC2A12** | 9.03E+01 | 3.64E-05 | | **IL36B** | -545.837 | 2.95E-05 |
| **SOX9-AS1** | 88.8054 | 0.000373 | | **KRT2** | -525.094 | 0.003388 |
| **SYNJ2BP-COX16** | 8.64E+01 | 0.000334 | | **BPIFC** | -473.749 | 3.85E-05 |
| **IL2RB** | 8.42E+01 | 4.69E-06 | | **SPRR2B** | -443.017 | 4.49E-05 |
| **LINC00479** | 8.04E+01 | 0.001093 | | **PSAPL1** | -417.493 | 0.000241 |
| **LOC101928053** | 80.0342 | 3.68E-05 | | **LCE1A** | -394.45 | 0.000207 |
| **LGI4** | 7.96E+01 | 6.17E-05 | | **KRT10** | -390.83 | 0.000119 |
| **LINC01376** | 7.81E+01 | 0.000193 | | **HOPX** | -384.553 | 4.60E-06 |
| **EFCAB5** | 7.63E+01 | 1.02E-06 | | **DHRS9** | -383.239 | 1.11E-05 |
| **KMO** | 74.9008 | 0.000163 | | **PTGER3** | -345.517 | 0.000467 |
| **FILIP1** | 71.6186 | 0.000474 | | **CLDN17** | -325.475 | 4.30E-05 |

| | | | | | | |
|---|---|---|---|---|---|---|
| **DINOL** | 7.11E+01 | 0.00016 | | **SPRR2E** | -307.866 | 3.42E-05 |
| **FAM198B-AS1** | 66.5646 | 1.08E-05 | | **RHBG** | -285.793 | 2.23E-05 |
| **VTCN1** | 63.6663 | 0.00082 | | **KRT4** | -276.277 | 3.83E-05 |
| **SERPINA3** | 61.0487 | 0.000662 | | **TGM5** | -266.141 | 5.05E-06 |
| **ITGB2** | 60.5344 | 0.00177 | | **ATP6V1C2** | -244.712 | 1.20E-06 |
| **PARM1** | 59.0635 | 0.000244 | | **ABCG4** | -242.96 | 4.59E-07 |
| **CLTRN** | 5.79E+01 | 2.77E-05 | | **TNFAIP8L3** | -242.543 | 6.20E-06 |
| **AOC1** | 5.71E+01 | 0.002126 | | **SLC26A9** | -235.507 | 4.11E-05 |
| **LOC100507516** | 5.64E+01 | 0.000117 | | **ACER1** | -226.151 | 1.48E-05 |
| **DNAH6** | 5.58E+01 | 1.66E-05 | | **PLA2G4E** | -225.191 | 1.06E-05 |
| **NBPF18P** | 54.0629 | 0.000334 | | **CHRNA9** | -221.967 | 0.001163 |
| **MEIG1** | 53.4328 | 1.57E-05 | | **PLA2G4D** | -221.691 | 2.08E-05 |
| **WFDC3** | 52.6211 | 5.04E-05 | | **EPHB6** | -203.678 | 8.31E-05 |
| **EPCAM** | 52.2977 | 9.04E-06 | | **WFDC5** | -201.612 | 2.30E-06 |
| **TBC1D8-AS1** | 51.3539 | 0.001522 | | **KRT6C** | -200.186 | 3.07E-05 |
| **COL11A2** | 50.8179 | 0.0001 | | **LCE3E** | -196.97 | 0.000177 |
| **MST1L** | 50.4896 | 1.09E-06 | | **MUCL1** | -193.82 | 0.00107 |
| **LMOD1** | 4.99E+01 | 0.000817 | | **MAL** | -189.973 | 0.000753 |
| **PRRT2** | 49.7234 | 0.003713 | | **PRR9** | -189.716 | 0.000353 |
| **NPY1R** | 48.2172 | 0.001092 | | **RDH12** | -182.282 | 2.75E-06 |
| **LINC01559** | 4.81E+01 | 0.00118 | | **SERPINB3** | -171.492 | 3.44E-06 |
| **TSPAN1** | 4.79E+01 | 0.000581 | | **WFDC12** | -165.542 | 0.000133 |
| **NR4A2** | 47.9064 | 0.000352 | | **LCE1B** | -164.126 | 1.17E-05 |
| **DCST1** | 47.5656 | 0.002207 | | **LINC01181** | -164.106 | 6.41E-05 |
| **MGAT4EP** | 47.5409 | 0.000131 | | **SEPTIN5** | -160.249 | 3.85E-05 |
| **NAALADL2** | 4.59E+01 | 0.000182 | | **LCE1E** | -156.48 | 5.52E-06 |
| **FAM72D** | 4.59E+01 | 0.000227 | | **FLG** | -156.399 | 0.000127 |
| **LOC101927245** | 45.6495 | 0.000244 | | **KRT77** | -154.956 | 0.000503 |
| **NALT1** | 4.45E+01 | 7.81E-05 | | **DSG1** | -147.467 | 3.27E-05 |
| **WFDC2** | 4.31E+01 | 0.003222 | | **TGM1** | -142.147 | 8.08E-06 |
| **MYOM2** | 41.0969 | 0.003568 | | **PGLYRP2** | -132.24 | 0.001665 |
| **H3C15** | 40.7531 | 0.001733 | | **SPTSSB** | -124.324 | 7.39E-07 |
| **H3C14** | 40.7531 | 0.001733 | | **VSIG8** | -123.858 | 0.00088 |
| **HCP5** | 40.5868 | 0.001113 | | **INSYN1** | -121.913 | 2.07E-05 |
| **GASK1B** | 39.8221 | 6.82E-05 | | **SLC5A1** | -121.406 | 3.10E-06 |
| **LINC00672** | 39.6686 | 0.000536 | | **SLC19A3** | -120.793 | 0.000874 |
| **LOC101929470** | 39.4324 | 7.37E-05 | | **ABHD12B** | -120.55 | 0.000362 |
| **LOC400710** | 39.2225 | 4.59E-05 | | **DGAT2** | -120.082 | 0.000192 |
| **THRB-AS1** | 39.1395 | 0.000424 | | **UGT1A7** | -115.332 | 0.000412 |
| **KCNN4** | 38.9807 | 9.01E-05 | | **TMEM170B** | -113.796 | 0.001942 |
| **HRK** | 38.9172 | 0.000773 | | **STXBP6** | -113.574 | 0.001018 |
| **CCR10** | 38.7024 | 0.001154 | | **ALDH5A1** | -111.29 | 3.17E-05 |
| **YBX2** | 3.84E+01 | 0.000205 | | **SLURP2** | -110.85 | 0.003163 |
| **SH2D6** | 3.79E+01 | 0.000791 | | **IL37** | -109.391 | 0.000575 |
| **VAMP1** | 3.77E+01 | 0.001482 | | **CALB2** | -106.635 | 0.000115 |
| **SERINC4** | 35.4103 | 2.28E-05 | | **H19** | -105.162 | 0.000128 |

| | | | | | | |
|---|---|---|---|---|---|---|
| ABHD11 | 35.1805 | 1.38E-05 | | KRT16P2 | -100.734 | 0.001823 |
| IRAIN | 3.52E+01 | 0.000185 | | TREX2 | -99.8615 | 1.88E-05 |
| LTBP3 | 34.8867 | 4.20E-06 | | PPP2R2C | -95.978 | 6.29E-05 |
| LOC101928617 | 34.5002 | 1.49E-05 | | PNPLA1 | -94.7267 | 0.000533 |
| TCHHL1 | 34.2884 | 0.00355 | | EVPLL | -93.0349 | 6.41E-05 |
| SLC9A5 | 34.1122 | 0.001226 | | LYPD2 | -91.4008 | 0.00093 |
| LOC101926889 | 33.9944 | 0.000754 | | ATP2C2 | -91.2726 | 2.25E-05 |
| NCEH1 | 33.7278 | 9.45E-07 | | SDR9C7 | -90.1501 | 0.000159 |
| FGR | 33.5631 | 0.000254 | | GJB6 | -88.2415 | 8.12E-05 |
| HLA-F-AS1 | 33.5257 | 0.003453 | | LCE3D | -86.8265 | 0.001885 |
| LINC02014 | 33.3705 | 1.49E-05 | | VSIG10L | -86.7942 | 0.000133 |
| PYHIN1 | 3.29E+01 | 0.000687 | | GPR78 | -86.4269 | 0.000397 |
| EDA2R | 32.5489 | 3.47E-07 | | C10orf99 | -82.9335 | 0.000125 |
| SNAI1 | 3.18E+01 | 0.001217 | | FAM25A | -81.2182 | 9.30E-06 |
| GRIN2C | 31.676 | 0.001138 | | RPTN | -77.5068 | 0.00015 |
| LOC100652758 | 31.6472 | 0.000517 | | CSTA | -77.4553 | 4.78E-07 |
| DYNLRB2 | 3.16E+01 | 0.001176 | | KLK1 | -76.8243 | 0.000125 |
| C9orf43 | 31.4849 | 0.002799 | | SPINK7 | -76.4101 | 3.52E-05 |
| HLA-F | 31.3991 | 0.001862 | | FABP5 | -74.5186 | 2.00E-05 |
| LEMD1 | 31.1807 | 0.000365 | | DSC1 | -73.8034 | 1.68E-05 |
| MICB | 30.7908 | 0.000269 | | KRT79 | -73.3875 | 0.001049 |
| CFAP43 | 30.6462 | 9.08E-05 | | LYPD5 | -73.0759 | 2.26E-05 |
| PRR15L | 3.06E+01 | 0.000668 | | IFNK | -72.6631 | 0.003563 |
| LINC02542 | 30.3633 | 5.51E-06 | | PAQR5 | -72.5225 | 0.00085 |
| LEKR1 | 2.97E+01 | 5.68E-05 | | SLC30A2 | -72.4475 | 0.001438 |
| ACVR2B-AS1 | 29.4369 | 5.15E-05 | | PTPN5 | -70.3418 | 0.001084 |
| FUCA1 | 29.0027 | 3.71E-07 | | ASPRV1 | -66.6115 | 0.000102 |
| ACTA2 | 2.89E+01 | 0.000273 | | KRT16P3 | -66.0653 | 0.000161 |
| CYP8B1 | 28.8544 | 0.000419 | | DEFB103A | -65.8061 | 0.000445 |
| GPD1 | 28.6247 | 0.000296 | | MUC22 | -65.5972 | 0.004202 |
| ZNF205-AS1 | 28.496 | 0.000165 | | CERS4 | -64.8481 | 6.98E-05 |
| CMTM7 | 28.3371 | 0.003744 | | KRT16P1 | -64.7372 | 0.000373 |
| DNAAF1 | 2.83E+01 | 0.000418 | | TAGLN3 | -63.6921 | 0.001213 |
| MMRN1 | 28.1351 | 3.50E-05 | | SPRR2A | -63.3649 | 0.000474 |
| TMEM225B | 2.81E+01 | 0.000153 | | ATP13A4 | -63.1139 | 6.45E-05 |
| C6orf52 | 27.9063 | 0.001095 | | LY6G6C | -61.8139 | 0.000275 |
| LINC01770 | 2.78E+01 | 0.000473 | | SLURP1 | -61.8034 | 0.002602 |
| PTPRH | 2.77E+01 | 0.004218 | | EPB41L3 | -61.6961 | 0.00127 |
| LINC01232 | 2.77E+01 | 0.000519 | | AQP9 | -61.5281 | 0.000593 |
| HLA-B | 27.3239 | 0.001372 | | PLA2G4E-AS1 | -60.1724 | 0.001629 |
| TOB2P1 | 2.71E+01 | 9.51E-09 | | ANKRD35 | -59.3124 | 2.15E-05 |
| FLJ31356 | 27.0245 | 0.000159 | | CRCT1 | -58.5339 | 0.000692 |
| EPS8 | 26.9058 | 2.05E-05 | | CALML5 | -58.0513 | 2.48E-05 |
| LOC100507053 | 2.69E+01 | 0.000199 | | PPP1R16B | -57.377 | 0.002206 |
| IL15 | 2.63E+01 | 0.000231 | | LINC02560 | -57.1664 | 3.57E-05 |
| ADCY4 | 26.3002 | 0.001394 | | SPRR2F | -57.0281 | 0.001395 |

| | | | | | | |
|---|---|---|---|---|---|---|
| **LINC01715** | 26.2097 | 0.00012 | | **FRMPD1** | -56.4857 | 0.003081 |
| **CDNF** | 2.58E+01 | 7.10E-05 | | **MYRF** | -55.2769 | 0.000169 |
| **RPL23AP87** | 25.714 | 7.46E-08 | | **UPK1A** | -54.6537 | 0.000448 |
| **UPK2** | 25.7078 | 0.003484 | | **CXCL14** | -54.6169 | 0.000421 |
| **OR10H1** | 25.6329 | 0.00031 | | **TMEM63C** | -54.3291 | 0.004383 |
| **LOC388242** | 2.56E+01 | 0.001083 | | **SLC15A1** | -54.3009 | 0.000212 |
| **LOC613038** | 25.6208 | 0.001083 | | **KRT78** | -53.2729 | 5.62E-05 |
| **LOC100289495** | 2.53E+01 | 0.000382 | | **METTL7A** | -52.909 | 0.001147 |
| **KLF3-AS1** | 25.2428 | 0.001185 | | **CPM** | -51.8402 | 4.34E-05 |
| **ARFGEF3** | 25.0281 | 6.18E-05 | | **CARMIL3** | -51.7249 | 0.000577 |
| **RHPN1-AS1** | 24.8736 | 0.000117 | | **CIDEA** | -51.5071 | 0.000158 |
| **FBXO15** | 2.48E+01 | 3.32E-05 | | **CYP4B1** | -51.2317 | 0.000133 |
| **MALINC1** | 24.803 | 6.79E-06 | | **IL1F10** | -50.4905 | 2.66E-06 |
| **CBX6** | 24.7976 | 5.64E-07 | | **SLC1A4** | -49.9968 | 0.000275 |
| **FAS** | 2.46E+01 | 6.66E-06 | | **LGALS7** | -48.8423 | 0.000162 |
| **GLIPR1L2** | 24.6295 | 0.000313 | | **S100A8** | -48.3323 | 2.72E-05 |
| **DNAJC27-AS1** | 24.4337 | 0.000495 | | **SPRR1A** | -47.8962 | 3.58E-05 |
| **LINC02535** | 24.4159 | 0.001792 | | **BBOX1** | -47.6586 | 0.000968 |
| **LINC00578** | 24.3732 | 0.001157 | | **SAMD5** | -46.6523 | 0.000503 |
| **SAMD12-AS1** | 24.1708 | 0.000432 | | **FAM89A** | -46.5968 | 6.55E-05 |
| **KLLN** | 24.1572 | 0.000471 | | **LGALS7B** | -46.5277 | 8.27E-06 |
| **TLR1** | 24.0886 | 3.15E-05 | | **GDF7** | -46.1984 | 0.000741 |
| **EFCAB12** | 2.40E+01 | 0.001066 | | **CRABP2** | -44.594 | 1.38E-05 |
| **LAMA2** | 23.9523 | 0.000336 | | **FCHO1** | -44.378 | 0.00022 |
| **LINC01732** | 23.7485 | 0.001684 | | **FABP4** | -43.1286 | 0.000662 |
| **LINC002481** | 2.36E+01 | 0.000152 | | **AKR1C2** | -42.7885 | 7.64E-06 |
| **CBLN3** | 2.34E+01 | 0.000209 | | **LINC00365** | -42.6171 | 0.002738 |
| **MAN1B1-DT** | 23.3646 | 0.000377 | | **BCL11A** | -42.5484 | 0.001478 |
| **ABCB9** | 23.3019 | 6.46E-06 | | **RAPGEF4** | -41.6753 | 0.000479 |
| **TMEM269** | 23.293 | 0.001671 | | **LINC01206** | -41.0898 | 0.0015 |
| **C2CD4A** | 23.2642 | 0.001507 | | **SPINK5** | -41.023 | 0.000355 |
| **LOC101928068** | 2.32E+01 | 0.000669 | | **NIPAL4** | -40.876 | 1.03E-05 |
| **LOC100288748** | 2.32E+01 | 1.30E-05 | | **SLC6A14** | -40.6733 | 1.55E-05 |
| **MICB-DT** | 2.30E+01 | 0.001302 | | **LINC01094** | -40.6512 | 6.99E-06 |
| **CXCL8** | 2.29E+01 | 1.13E-05 | | **USH1G** | -40.5298 | 0.001001 |
| **LINC00184** | 22.859 | 0.000342 | | **NPR2** | -40.3561 | 2.51E-05 |
| **ENKUR** | 2.28E+01 | 0.004182 | | **S1PR5** | -40.2393 | 0.001537 |
| **DPP4** | 2.28E+01 | 0.001245 | | **DLX5** | -40.1807 | 0.003865 |
| **CD163L1** | 22.4707 | 0.000356 | | **RASD2** | -39.9825 | 0.000145 |
| **TNKS2-AS1** | 22.4615 | 1.26E-05 | | **KRT6B** | -39.7676 | 1.38E-05 |
| **CASQ2** | 22.4305 | 0.003677 | | **LGALSL** | -39.6824 | 8.13E-06 |
| **ZNF488** | 22.3113 | 0.00042 | | **TMEM45A** | -39.6512 | 0.00023 |
| **METTL27** | 22.2183 | 0.003326 | | **XKRX** | -38.9749 | 0.00058 |
| **CD79A** | 21.9614 | 0.001301 | | **SERPINB4** | -38.584 | 1.63E-05 |
| **MUC16** | 21.9284 | 0.000712 | | **FRY** | -38.4026 | 1.23E-05 |
| **MUC19** | 21.8698 | 0.001423 | | **AIF1L** | -38.2571 | 4.46E-05 |

| | | | | | | |
|---|---|---|---|---|---|---|
| **SYT14** | 2.18E+01 | 0.001076 | | **AKR1C3** | -38.2362 | 3.28E-05 |
| **CLDN3** | 2.17E+01 | 0.001319 | | **KCTD15** | -38.1433 | 1.81E-05 |
| **IGSF11** | 21.667 | 0.002625 | | **SEPTIN3** | -37.9325 | 0.001962 |
| **NT5E** | 2.17E+01 | 0.00149 | | **GLRX** | -37.6633 | 0.000503 |
| **LOC101928728** | 21.4756 | 0.000898 | | **SLC6A2** | -37.3086 | 0.000205 |
| **TIPARP-AS1** | 2.11E+01 | 0.000454 | | **CNFN** | -37.1636 | 0.000293 |
| **LINC00519** | 21.078 | 0.00016 | | **CYBRD1** | -36.7883 | 9.42E-05 |
| **ECT2L** | 21.0369 | 2.13E-05 | | **SCNN1B** | -36.4512 | 0.001132 |
| **LRRC46** | 20.9909 | 0.000832 | | **RUNDC3A** | -36.1883 | 0.000612 |
| **PDE4C** | 2.10E+01 | 6.64E-06 | | **PALMD** | -36.0369 | 3.10E-05 |
| **BMP6** | 20.9219 | 1.47E-05 | | **NES** | -36.0304 | 5.40E-05 |
| **SLC15A3** | 20.8626 | 0.00435 | | **GFRA1** | -35.1391 | 0.004318 |
| **C19orf38** | 20.6846 | 0.000821 | | **NFATC2** | -35.1208 | 0.000101 |
| **HLA-H** | 20.5948 | 0.000498 | | **CEACAM7** | -34.5564 | 0.001859 |
| **GUCA1B** | 20.5367 | 2.55E-07 | | **CDHR1** | -34.4836 | 0.000384 |
| **SERTAD4** | 2.05E+01 | 0.002039 | | **LRRC75B** | -34.4636 | 0.003491 |
| **RRM2B** | 2.04E+01 | 1.80E-06 | | **AKR1C1** | -34.3686 | 2.61E-05 |
| **ID2-AS1** | 20.3472 | 8.51E-06 | | **ESYT3** | -34.3678 | 0.003822 |
| **NCMAP** | 2.03E+01 | 0.001325 | | **TENM2** | -34.0289 | 0.000113 |
| **PLA2G10** | 20.1412 | 0.000116 | | **PI3** | -33.8194 | 0.000227 |
| **IL15RA** | 19.9766 | 0.000171 | | **SERPINB11** | -33.4429 | 0.00025 |
| **MST1P2** | 19.933 | 0.000311 | | **ARRB1** | -33.3087 | 0.000219 |
| **SRRM2-AS1** | 1.98E+01 | 0.000506 | | **TMPRSS11A** | -33.2497 | 0.001025 |
| **CDRT4** | 1.97E+01 | 0.000551 | | **GNG4** | -33.1116 | 5.85E-05 |
| **CBX3P2** | 19.719 | 0.003183 | | **RNF225** | -32.7351 | 0.000912 |
| **RPGRIP1** | 19.6993 | 1.80E-05 | | **HCG22** | -32.4536 | 0.000101 |
| **PGF** | 1.97E+01 | 2.91E-05 | | **LINC01224** | -32.2591 | 0.001793 |
| **TNFRSF10D** | 19.4539 | 5.35E-05 | | **CD36** | -32.2342 | 0.000116 |
| **LINC00880** | 19.3389 | 0.000986 | | **SLC46A2** | -32.1786 | 0.000178 |
| **NPIPB6** | 19.3207 | 0.000262 | | **GJB2** | -31.8263 | 9.68E-06 |
| **SEC14L5** | 1.90E+01 | 2.16E-05 | | **PALD1** | -31.822 | 0.001493 |
| **ASIC3** | 18.4923 | 0.001197 | | **SPRR2D** | -31.6408 | 0.000583 |
| **SAP30L-AS1** | 1.85E+01 | 8.09E-05 | | **P2RY1** | -31.3837 | 0.000114 |
| **TRIM74** | 1.85E+01 | 0.0002 | | **ALOXE3** | -31.3064 | 0.000123 |
| **MAJIN** | 18.3051 | 0.001475 | | **IL36G** | -31.144 | 0.000278 |
| **ATF3** | 18.2874 | 0.00041 | | **RBP7** | -31.0879 | 0.000333 |
| **CATSPERG** | 1.82E+01 | 2.17E-05 | | **SULT1E1** | -30.7692 | 0.003637 |
| **GPR35** | 18.2118 | 0.000744 | | **S100A7** | -30.7562 | 0.000666 |
| **LINC01251** | 18.1988 | 0.000187 | | **TCEA3** | -30.3365 | 0.001859 |
| **KLF9** | 18.1503 | 3.72E-05 | | **TUBB2A** | -30.1885 | 9.13E-05 |
| **COL28A1** | 18.1225 | 0.00349 | | **APCDD1L-DT** | -30.111 | 0.00327 |
| **SEMA3E** | 1.81E+01 | 0.000528 | | **WFDC21P** | -30.0264 | 0.000312 |
| **LINC02029** | 18.0475 | 0.002018 | | **NMRAL2P** | -29.4914 | 5.45E-05 |
| **CDKL3** | 18.0372 | 0.001938 | | **GPRIN2** | -29.4883 | 0.000225 |
| **DLGAP1-AS2** | 18.015 | 0.00086 | | **CHRNA3** | -29.4083 | 0.001853 |
| **LINC00449** | 1.80E+01 | 0.000866 | | **TMPRSS11B** | -29.0759 | 0.002348 |

| | | | | | | |
|---|---|---|---|---|---|---|
| **CDKN2B-AS1** | 17.7674 | 3.91E-06 | | **HMOX1** | -28.9422 | 0.000152 |
| **BDNF-AS** | 17.751 | 0.000754 | | **LOC101929331** | -28.8643 | 0.000148 |
| **FAM81A** | 17.7225 | 0.000883 | | **PLXDC2** | -28.8439 | 0.000144 |
| **ITLN2** | 1.77E+01 | 0.001337 | | **ACSBG1** | -28.8192 | 0.000479 |
| **MYO15A** | 1.77E+01 | 4.67E-05 | | **RASAL1** | -28.7014 | 6.06E-05 |
| **TENT5A** | 17.69 | 7.02E-06 | | **FCHSD1** | -28.5775 | 7.03E-05 |
| **LINC01588** | 17.6475 | 9.94E-06 | | **LCE5A** | -28.3042 | 0.002822 |
| **MEP1A** | 17.6047 | 0.00222 | | **ANKFN1** | -28.1955 | 0.00138 |
| **ACKR2** | 17.5409 | 0.000237 | | **HOMER2** | -28.0388 | 1.26E-05 |
| **FER1L6** | 17.4836 | 0.000283 | | **FOXN1** | -27.7895 | 0.000409 |
| **HOXA-AS2** | 17.4488 | 4.57E-06 | | **SLC29A4** | -27.3732 | 0.000176 |
| **IRF1** | 17.4044 | 0.00018 | | **ACP7** | -27.3386 | 6.99E-05 |
| **BIK** | 17.3376 | 9.73E-05 | | **APOL4** | -27.1694 | 0.000159 |
| **BAALC-AS1** | 17.1093 | 0.000165 | | **BOC** | -27.147 | 0.002664 |
| **ERAP2** | 17.0219 | 5.43E-05 | | **CAPNS2** | -27.032 | 5.99E-06 |
| **SULF2** | 16.9927 | 5.60E-05 | | **CCDC8** | -26.9583 | 0.000343 |
| **PAPPA** | 1.68E+01 | 2.44E-05 | | **BLMH** | -26.8979 | 0.000444 |
| **HOTAIR** | 16.8388 | 0.000845 | | **RHCG** | -26.3406 | 7.71E-06 |
| **ASB9** | 16.8301 | 0.001737 | | **APOB** | -26.2884 | 0.003727 |
| **SLAMF9** | 1.68E+01 | 0.001013 | | **SPRR1B** | -26.2362 | 3.00E-05 |
| **PCNA** | 16.7723 | 2.27E-05 | | **GLB1L2** | -26.1464 | 0.000308 |
| **DRC3** | 16.7514 | 0.000133 | | **FOXP2** | -25.9742 | 0.001622 |
| **RPSAP52** | 16.7319 | 0.00436 | | **MAB21L3** | -25.9271 | 4.70E-05 |
| **NTN4** | 1.67E+01 | 1.38E-06 | | **TMEM37** | -25.888 | 0.003914 |
| **LOC102723665** | 1.66E+01 | 0.00121 | | **SLC7A4** | -25.8156 | 2.57E-05 |
| **AQP7** | 1.66E+01 | 0.001748 | | **SPRR4** | -25.7966 | 0.000432 |
| **CXCL6** | 16.588 | 0.000287 | | **SERPINB12** | -25.7414 | 6.47E-05 |
| **CAMK2N1** | 16.4511 | 7.73E-05 | | **CCDC3** | -25.706 | 0.00029 |
| **LOC101929762** | 16.366 | 0.001803 | | **ARHGAP40** | -25.6736 | 4.60E-05 |
| **CA3-AS1** | 1.63E+01 | 0.000147 | | **KIF26B** | -25.673 | 0.000531 |
| **LOC100507634** | 1.63E+01 | 0.000889 | | **LINC02159** | -25.4889 | 0.000166 |
| **SLC35G6** | 16.2495 | 0.001038 | | **S100A9** | -25.4578 | 0.000199 |
| **KLRD1** | 1.62E+01 | 0.000307 | | **INSYN2B** | -25.343 | 5.21E-05 |
| **PRKG1-AS1** | 16.2042 | 0.001143 | | **KRT75** | -25.2375 | 0.002886 |
| **SSPN** | 16.1737 | 5.11E-05 | | **TMEM238L** | -25.2038 | 0.001125 |
| **TSGA10** | 1.61E+01 | 1.11E-05 | | **CRYAB** | -24.5157 | 0.000599 |
| **SYCE2** | 16.0931 | 0.000744 | | **RASGRP1** | -24.4621 | 0.000213 |
| **LINC00663** | 1.60E+01 | 0.000249 | | **SMPD3** | -24.334 | 0.001841 |
| **LINC00886** | 16.0138 | 0.000238 | | **EFR3B** | -24.3134 | 0.000327 |
| **TF** | 1.60E+01 | 0.004409 | | **SULT2B1** | -24.2517 | 3.83E-05 |
| **CRACR2B** | 15.9803 | 0.000259 | | **ASAP3** | -24.1308 | 0.000279 |
| **REM2** | 15.8977 | 0.000442 | | **KRT6A** | -24.1017 | 1.21E-05 |
| **LOC101927911** | 15.888 | 3.72E-05 | | **AKR1B10** | -24.0449 | 0.000382 |
| **LINC00342** | 1.58E+01 | 0.000623 | | **MECOM-AS1** | -24.027 | 0.002871 |
| **LOC644285** | 15.826 | 0.001233 | | **PHGDH** | -23.98 | 0.000157 |
| **TRHDE** | 15.802 | 0.001202 | | **ANXA9** | -23.8827 | 1.69E-08 |

| | | | | | | |
|---|---|---|---|---|---|---|
| **HLA-C** | 1.57E+01 | 0.000261 | | **HS3ST6** | -23.393 | 0.000553 |
| **LOC101927811** | 15.7213 | 0.000128 | | **FETUB** | -23.1706 | 0.000262 |
| **COMP** | 1.57E+01 | 2.17E-05 | | **KRTDAP** | -23.0103 | 0.000122 |
| **RARB** | 1.56E+01 | 0.001553 | | **CYP39A1** | -22.7701 | 2.05E-05 |
| **STKLD1** | 1.56E+01 | 0.001809 | | **PCSK9** | -22.6956 | 0.001051 |
| **CLDN23** | 15.5689 | 3.21E-06 | | **AZGP1** | -22.6587 | 0.002656 |
| **SFR1** | 15.5143 | 1.30E-05 | | **PNLIPRP3** | -22.6296 | 2.87E-05 |
| **RASGRF1** | 1.55E+01 | 3.80E-05 | | **GGT8P** | -22.5556 | 1.84E-05 |
| **ABCD1** | 15.4581 | 0.001366 | | **NLRP10** | -22.5153 | 0.001914 |
| **WNK4** | 15.4063 | 0.001888 | | **ELF5** | -22.4019 | 0.000104 |
| **MAP3K14-AS1** | 1.54E+01 | 9.87E-06 | | **BICC1** | -21.9387 | 0.000153 |
| **UBR5-AS1** | 1.53E+01 | 0.000533 | | **C12orf56** | -21.6361 | 0.00143 |
| **B2M** | 1.52E+01 | 0.001402 | | **USP2** | -21.5823 | 0.000181 |
| **RAD51AP2** | 1.52E+01 | 0.001805 | | **GPSM1** | -21.4418 | 0.00045 |
| **NEDD9** | 1.52E+01 | 7.63E-05 | | **SPRR3** | -21.397 | 0.000201 |
| **FCGBP** | 15.1477 | 5.25E-05 | | **GFOD1** | -21.3782 | 1.85E-05 |
| **RBM26-AS1** | 15.0437 | 0.000331 | | **GLTP** | -21.3556 | 1.84E-06 |
| **ARL14** | 1.50E+01 | 0.000689 | | **SERPINB13** | -21.2267 | 3.31E-06 |
| **KRT8** | 14.9936 | 0.000298 | | **LRRC20** | -21.1833 | 2.80E-05 |
| **NKAPL** | 1.50E+01 | 0.00134 | | **ADTRP** | -20.9546 | 1.97E-05 |
| **LOC107986163** | 1.49E+01 | 0.000361 | | **PLBD1-AS1** | -20.93 | 0.000437 |
| **BHLHE40-AS1** | 1.48E+01 | 0.00027 | | **GJA1** | -20.8356 | 2.71E-05 |
| **FLJ37453** | 1.48E+01 | 0.000797 | | **RAPGEFL1** | -20.8083 | 1.27E-05 |
| **ANKRD36C** | 1.47E+01 | 7.18E-05 | | **ASIC1** | -20.7087 | 0.003606 |
| **LINC00857** | 14.7081 | 2.60E-05 | | **NUPR1** | -20.7017 | 0.000149 |
| **P4HTM** | 1.47E+01 | 1.89E-05 | | **LINC01322** | -20.692 | 0.001415 |
| **HEXA-AS1** | 14.6208 | 0.001898 | | **ENDOU** | -20.5807 | 1.91E-05 |
| **TAS1R3** | 14.6201 | 0.001983 | | **ALOX15B** | -20.525 | 0.000857 |
| **GSN-AS1** | 14.4793 | 0.003833 | | **HKDC1** | -20.4395 | 0.00026 |
| **LINC01679** | 14.4567 | 0.000111 | | **ELOVL3** | -20.4071 | 0.004216 |
| **C1QTNF1** | 1.44E+01 | 0.000419 | | **KLK12** | -20.3622 | 0.000243 |
| **LRRC4** | 14.2893 | 0.00257 | | **SQLE** | -20.2723 | 7.66E-05 |
| **DUSP4** | 14.2507 | 0.001163 | | **KLK14** | -20.2395 | 0.000891 |
| **NPIPB2** | 14.2504 | 0.000294 | | **NOD2** | -20.2335 | 0.000146 |
| **MB** | 1.42E+01 | 0.000486 | | **ZNF662** | -20.205 | 0.000125 |
| **STEAP3** | 14.1694 | 2.16E-06 | | **S100A12** | -20.0583 | 0.003071 |
| **TNFSF9** | 1.41E+01 | 3.38E-05 | | **IL33** | -20.046 | 0.003525 |
| **SPATA18** | 14.1135 | 7.15E-07 | | **SLC47A1** | -20.002 | 0.003204 |
| **LOC112543491** | 13.9234 | 0.001431 | | **SLC16A5** | -19.9063 | 0.004317 |
| **PDCD4-AS1** | 13.9053 | 0.002102 | | **GAS7** | -19.9036 | 0.000394 |
| **CNTNAP1** | 1.39E+01 | 0.00309 | | **CYP4F12** | -19.7475 | 0.001216 |
| **LOC100288798** | 13.855 | 6.90E-05 | | **LAMB4** | -19.6917 | 0.001773 |
| **CD70** | 1.39E+01 | 0.001085 | | **THEM5** | -19.6758 | 4.32E-05 |
| **LOC101928504** | 13.8044 | 0.000104 | | **EGR2** | -19.6657 | 7.85E-05 |
| **DKK1** | 1.37E+01 | 0.000346 | | **SDK2** | -19.6502 | 2.70E-05 |
| **TONSL-AS1** | 13.7102 | 0.000975 | | **CSPG5** | -19.5461 | 0.004169 |

| | | | | | | |
|---|---|---|---|---|---|---|
| **ABCC11** | 13.701 | 0.000759 | | **SEMA4D** | -19.4934 | 2.05E-05 |
| **C11orf91** | 1.37E+01 | 0.000177 | | **CMTM8** | -19.326 | 0.001623 |
| **ZNF702P** | 13.6549 | 0.002768 | | **FAM83C** | -19.3242 | 8.49E-05 |
| **PLS1** | 1.36E+01 | 9.69E-05 | | **PSORS1C2** | -19.3068 | 7.93E-05 |
| **DSG2** | 1.36E+01 | 2.23E-05 | | **PLA2G4B** | -19.2938 | 2.75E-05 |
| **PPP1R26-AS1** | 13.6245 | 0.000379 | | **SMOX** | -19.1855 | 2.34E-05 |
| **GDF15** | 13.5911 | 6.84E-05 | | **RIMS3** | -19.173 | 0.000215 |
| **TSPAN15** | 1.36E+01 | 1.24E-05 | | **DLL1** | -19.1007 | 0.000717 |
| **DNAH10** | 13.5559 | 0.0011 | | **PDE9A** | -18.8595 | 0.001672 |
| **DEPP1** | 1.35E+01 | 0.002557 | | **CNKSR3** | -18.8099 | 4.46E-05 |
| **HLA-J** | 13.464 | 0.000846 | | **NXPH3** | -18.7303 | 0.000119 |
| **LINC00638** | 1.34E+01 | 0.000641 | | **IL36RN** | -18.6698 | 5.29E-05 |
| **LINC01451** | 13.4217 | 0.001475 | | **LRP4** | -18.4512 | 0.001618 |
| **FYB1** | 13.3639 | 0.000397 | | **SNPH** | -18.4503 | 0.000585 |
| **UBAC2-AS1** | 1.34E+01 | 0.000248 | | **LINC01133** | -18.3764 | 2.00E-05 |
| **FHAD1** | 13.3177 | 9.48E-05 | | **KRT16** | -18.1644 | 0.000232 |
| **NEU1** | 13.2161 | 4.78E-05 | | **ACOT11** | -18.0481 | 2.97E-05 |
| **SERPINB10** | 1.32E+01 | 0.000143 | | **TENT5B** | -18.0241 | 8.78E-05 |
| **LOC105371763** | 1.32E+01 | 0.000636 | | **SYNGR1** | -18.0229 | 0.000396 |
| **NTN5** | 1.32E+01 | 0.003622 | | **MIR205HG** | -17.9688 | 7.93E-07 |
| **STAG3** | 1.31E+01 | 0.001188 | | **FAM83D** | -17.9578 | 0.000291 |
| **CXCL1** | 13.063 | 0.000831 | | **IGFL2** | -17.9262 | 1.12E-06 |
| **TG** | 1.31E+01 | 0.000545 | | **SLC9A9** | -17.9002 | 0.000181 |
| **SPRED3** | 13.0123 | 5.68E-05 | | **TNNT1** | -17.8159 | 3.97E-05 |
| **CRPPA** | 12.9991 | 8.81E-05 | | **GSDMA** | -17.6847 | 0.000773 |
| **CEL** | 12.9835 | 5.09E-06 | | **NPAS1** | -17.394 | 0.000601 |
| **NLRC5** | 12.8319 | 0.000828 | | **IGFL3** | -17.3737 | 0.000355 |
| **RPL13AP20** | 1.28E+01 | 0.000275 | | **OLFM2** | -17.3019 | 6.96E-06 |
| **CA5BP1-CA5B** | 1.28E+01 | 6.07E-06 | | **NGEF** | -17.2709 | 0.001095 |
| **C16orf71** | 12.8201 | 4.67E-05 | | **TCAF2** | -17.2086 | 0.001518 |
| **MICA** | 12.816 | 0.000124 | | **RASSF9** | -17.1682 | 0.000287 |
| **CCN4** | 12.8027 | 2.53E-05 | | **EDDM13** | -17.1165 | 9.88E-05 |
| **ST3GAL4** | 12.757 | 1.38E-05 | | **COL8A2** | -16.8591 | 0.003613 |
| **LINC01909** | 12.7261 | 0.002897 | | **JDP2** | -16.7995 | 0.000302 |
| **HLA-L** | 12.6608 | 0.001501 | | **KRT23** | -16.7951 | 0.000312 |
| **ARSB** | 12.6293 | 0.000352 | | **A2ML1** | -16.794 | 5.23E-05 |
| **HERC5** | 1.26E+01 | 0.003097 | | **KCNS1** | -16.7928 | 0.000153 |
| **EIF2AK3-DT** | 12.5755 | 9.56E-05 | | **IL27RA** | -16.7926 | 0.001127 |
| **DUSP28** | 12.5655 | 0.0001 | | **ALDH3B2** | -16.7871 | 1.40E-05 |
| **LOC101929268** | 1.25E+01 | 0.000803 | | **NT5DC2** | -16.7613 | 2.54E-05 |
| **LOC100507564** | 12.4938 | 7.44E-05 | | **SLC13A4** | -16.7104 | 0.000257 |
| **PTGES** | 1.25E+01 | 0.001088 | | **CAMK1D** | -16.5922 | 3.22E-05 |
| **GALNT7** | 1.24E+01 | 2.29E-05 | | **SLC22A3** | -16.5155 | 0.000328 |
| **CATSPERB** | 1.23E+01 | 0.003888 | | **NDUFA4L2** | -16.3814 | 0.003849 |
| **BEST3** | 12.3144 | 0.002024 | | **KRT17** | -16.3587 | 3.38E-05 |
| **CCT6B** | 12.2218 | 8.28E-05 | | **IL20RB** | -16.3505 | 0.000107 |

| | | | | | | |
|---|---|---|---|---|---|---|
| **KIAA0040** | 12.1805 | 0.000107 | | **RAB3D** | -16.2629 | 1.68E-05 |
| **SLC4A9** | 12.1699 | 0.000166 | | **CWH43** | -16.2301 | 0.000153 |
| **LOC645967** | 12.1646 | 0.000276 | | **ME1** | -15.9786 | 5.24E-05 |
| **LOC283045** | 1.22E+01 | 0.00056 | | **KRT14** | -15.8787 | 0.000182 |
| **CILP2** | 12.1584 | 0.001614 | | **ACP3** | -15.8752 | 2.95E-05 |
| **LINC00942** | 12.1333 | 0.001112 | | **SHISAL1** | -15.8207 | 0.001157 |
| **NKILA** | 12.131 | 8.21E-06 | | **SEMA6D** | -15.7785 | 0.003238 |
| **TLE6** | 12.0979 | 0.000413 | | **SMO** | -15.7515 | 5.85E-05 |
| **BCL2L11** | 12.0888 | 0.00011 | | **MTSS1** | -15.6939 | 3.28E-06 |
| **ACER2** | 12.0269 | 1.56E-05 | | **SLC16A6** | -15.6285 | 0.000388 |
| **CASP7** | 11.9842 | 3.45E-06 | | **SGPP2** | -15.6182 | 1.89E-05 |
| **DHX58** | 11.9796 | 0.000108 | | **TRIM7** | -15.5999 | 0.000113 |
| **LINC02580** | 11.967 | 5.48E-05 | | **CYP4F11** | -15.5573 | 0.001267 |
| **ZNF341-AS1** | 11.9321 | 1.34E-05 | | **CFAP58-DT** | -15.5448 | 0.001086 |
| **CREG2** | 11.8268 | 0.001828 | | **CD248** | -15.4842 | 0.001527 |
| **SEC31B** | 11.8233 | 0.001814 | | **FMO2** | -15.4595 | 0.000717 |
| **ABCA9** | 11.8088 | 0.000544 | | **HYAL1** | -15.4279 | 6.67E-07 |
| **C2orf92** | 11.7888 | 0.003595 | | **TNK2-AS1** | -15.4147 | 0.003175 |
| **PPIL6** | 11.7296 | 0.000233 | | **KCNB1** | -15.2613 | 0.000527 |
| **APH1B** | 11.6875 | 0.000144 | | **ABLIM1** | -15.2423 | 2.24E-05 |
| **GLB1L** | 1.17E+01 | 2.62E-06 | | **PGLYRP3** | -15.2218 | 5.75E-05 |
| **OSR2** | 1.17E+01 | 0.000262 | | **GDPD2** | -15.1985 | 0.001469 |
| **AKAP3** | 11.6547 | 8.64E-06 | | **FAM43A** | -15.1838 | 6.37E-05 |
| **SMOC1** | 1.16E+01 | 0.000992 | | **LARGE2** | -15.0714 | 0.000801 |
| **NPBWR1** | 11.6174 | 0.003422 | | **PCK2** | -15.0584 | 0.000603 |
| **C1GALT1C1** | 11.6115 | 1.97E-05 | | **ADAP2** | -15.026 | 1.11E-05 |
| **LOC100287329** | 11.595 | 0.001713 | | **AMZ1** | -14.9875 | 0.000227 |
| **ARID3C** | 11.5624 | 2.74E-05 | | **SLC47A2** | -14.9721 | 0.000542 |
| **RASGRP3** | 1.15E+01 | 0.002008 | | **SBSN** | -14.9372 | 0.001532 |
| **ANKDD1B** | 1.15E+01 | 0.000197 | | **ATP10B** | -14.7972 | 0.00038 |
| **RSPH10B2** | 1.15E+01 | 0.000632 | | **SERPINB9** | -14.7556 | 0.000222 |
| **RSPH10B** | 11.4761 | 0.000632 | | **TMEM86A** | -14.7307 | 0.000554 |
| **LINC00899** | 11.456 | 0.0003 | | **SORD** | -14.649 | 1.16E-05 |
| **IRF1-AS1** | 11.454 | 0.000611 | | **APCDD1** | -14.533 | 0.001096 |
| **PSPN** | 1.14E+01 | 4.06E-05 | | **PCSK6** | -14.4852 | 4.82E-05 |
| **TAF1A-AS1** | 11.4299 | 0.000156 | | **RAET1E** | -14.4522 | 3.99E-05 |
| **LOC646626** | 11.4269 | 2.85E-05 | | **ZDHHC23** | -14.4447 | 0.001771 |
| **HTR7P1** | 11.4111 | 4.16E-05 | | **PLA2G4F** | -14.4377 | 7.91E-06 |
| **FAM107B** | 11.4022 | 6.50E-05 | | **CA12** | -14.3969 | 3.30E-05 |
| **LMNTD2** | 1.14E+01 | 0.000201 | | **ACBD3-AS1** | -14.3753 | 0.004268 |
| **ANKRD2** | 11.3771 | 0.002614 | | **RTN4RL1** | -14.2856 | 0.001569 |
| **GLIS3** | 11.3597 | 0.002082 | | **NSG1** | -14.2383 | 9.59E-05 |
| **MGAM** | 1.13E+01 | 0.00078 | | **OVCH2** | -14.1878 | 3.71E-06 |
| **LOC101928069** | 11.3202 | 0.000112 | | **EEPD1** | -14.1547 | 0.001919 |
| **SNX29P1** | 1.13E+01 | 0.002085 | | **PGAP4** | -14.0395 | 6.16E-05 |
| **AOC3** | 11.3028 | 0.000258 | | **CERS3** | -14.0084 | 4.27E-05 |

| | | | | | | |
|---|---|---|---|---|---|---|
| **PSMB9** | 1.13E+01 | 0.000532 | | **CLIC3** | -13.9942 | 0.000586 |
| **C1orf105** | 11.2751 | 0.000235 | | **LINC01214** | -13.9782 | 0.001192 |
| **LAMP3** | 11.2042 | 0.000296 | | **ACKR3** | -13.8801 | 0.000555 |
| **PCBP1-AS1** | 1.12E+01 | 0.00021 | | **SUSD4** | -13.7813 | 3.88E-05 |
| **WNT9A** | 1.12E+01 | 0.00228 | | **PLA2R1** | -13.7804 | 1.30E-05 |
| **MYPN** | 11.15 | 0.000232 | | **SLC7A11** | -13.7276 | 0.000965 |
| **TRHDE-AS1** | 11.1249 | 0.002586 | | **GBP6** | -13.5733 | 0.000149 |
| **ABHD11-AS1** | 11.1173 | 0.002478 | | **NKPD1** | -13.5621 | 0.000214 |
| **ACBD7** | 11.0643 | 0.00078 | | **NGFR** | -13.5603 | 0.002746 |
| **LINC00205** | 11.0416 | 6.69E-05 | | **QPRT** | -13.5353 | 9.36E-05 |
| **HLA-A** | 11.0331 | 0.002087 | | **DNASE1L3** | -13.5268 | 5.40E-06 |
| **RHOU** | 1.10E+01 | 0.002527 | | **ZHX3** | -13.4971 | 0.000166 |
| **ARHGAP5-AS1** | 10.9962 | 0.000217 | | **FAAHP1** | -13.454 | 0.001478 |
| **NPC1L1** | 10.9772 | 0.000922 | | **OBSCN** | -13.447 | 4.02E-05 |
| **AOC2** | 10.9743 | 0.00023 | | **CBSL** | -13.4282 | 0.002442 |
| **GULP1** | 1.10E+01 | 8.38E-05 | | **VWA2** | -13.3771 | 0.003324 |
| **AQP7P3** | 10.967 | 0.001951 | | **ZC3HAV1L** | -13.3425 | 0.002054 |
| **LPP-AS2** | 1.10E+01 | 0.000233 | | **ATP12A** | -13.3187 | 0.001404 |
| **DNAH1** | 10.9424 | 0.002405 | | **CHST2** | -13.2997 | 0.002368 |
| **GALNT4** | 10.9339 | 0.000829 | | **FNDC10** | -13.2936 | 6.85E-05 |
| **MIR762HG** | 10.8881 | 0.000545 | | **DIAPH3** | -13.2264 | 0.000227 |
| **RGPD4** | 1.09E+01 | 0.001493 | | **RNF222** | -13.2251 | 0.000334 |
| **ERFL** | 10.8288 | 0.000563 | | **ST6GALNAC1** | -13.1832 | 8.80E-05 |
| **BSCL2** | 10.8151 | 0.000926 | | **COL27A1** | -13.1649 | 0.000932 |
| **APOL6** | 10.7806 | 1.03E-05 | | **ADM2** | -13.1522 | 0.002627 |
| **TAPBPL** | 10.7578 | 7.75E-05 | | **CDSN** | -13.1467 | 0.000245 |
| **WDR78** | 10.72 | 2.19E-05 | | **TTC39B** | -13.0998 | 5.17E-05 |
| **RORA-AS1** | 10.6569 | 0.002542 | | **FAT2** | -12.9287 | 1.52E-06 |
| **REPS2** | 10.6393 | 0.000324 | | **ALDH2** | -12.9155 | 0.000994 |
| **BFSP1** | 1.06E+01 | 0.000637 | | **ASS1** | -12.9148 | 0.001501 |
| **LINC01465** | 1.06E+01 | 0.00214 | | **BMPER** | -12.8677 | 0.001859 |
| **PLK5** | 10.5945 | 0.000139 | | **TTTY4C** | -12.8655 | 0.000135 |
| **TBC1D8** | 10.5758 | 7.48E-06 | | **TTTY4B** | -12.8655 | 0.000135 |
| **CDKN2A** | 10.5708 | 0.000245 | | **TTTY4** | -12.8655 | 0.000135 |
| **VEGFC** | 10.5612 | 5.16E-05 | | **DNMT3B** | -12.8419 | 0.000297 |
| **TOLLIP-AS1** | 10.5591 | 0.000759 | | **CDCA7** | -12.8375 | 0.000152 |
| **PODXL** | 10.5367 | 0.003104 | | **WNT4** | -12.8348 | 0.000486 |
| **TRAK2** | 1.05E+01 | 0.001444 | | **ZBTB7C** | -12.7909 | 3.32E-05 |
| **H4C15** | 10.4501 | 0.000225 | | **FHL1** | -12.7518 | 0.000602 |
| **H4C14** | 10.4501 | 0.000225 | | **GSTA4** | -12.691 | 1.76E-06 |
| **STAT4** | 1.04E+01 | 0.000103 | | **GGCT** | -12.5198 | 0.00014 |
| **TMPRSS4** | 1.04E+01 | 8.57E-05 | | **IGFL1** | -12.4606 | 0.001061 |
| **ANGPT2** | 1.04E+01 | 0.003865 | | **MFSD2B** | -12.3972 | 0.0005 |
| **LOC102723769** | 1.04E+01 | 0.00199 | | **VWF** | -12.3672 | 8.48E-05 |
| **ADAM15** | 10.3671 | 2.40E-05 | | **NPM2** | -12.301 | 0.003102 |
| **ZNF528-AS1** | 10.3101 | 0.000403 | | **SLC10A6** | -12.243 | 0.00031 |

| | | | | | | |
|---|---|---|---|---|---|---|
| **CCDC87** | 1.03E+01 | 0.000101 | | **FOXE1** | -12.2391 | 1.81E-05 |
| **CYP2C8** | 10.2924 | 0.003444 | | **FGFBP1** | -12.1867 | 0.000871 |
| **LINC02298** | 10.2554 | 0.00103 | | **TMTC1** | -12.1462 | 0.000336 |
| **PXDNL** | 10.2336 | 0.000444 | | **ARL10** | -12.1435 | 1.15E-06 |
| **LINC01415** | 10.2217 | 0.003707 | | **EHD2** | -12.1 | 0.000705 |
| **LINC01607** | 10.1897 | 0.001475 | | **PLA2G2F** | -12.0905 | 0.000434 |
| **TCTN3** | 10.178 | 7.29E-06 | | **HMCN1** | -12.0603 | 0.00154 |
| **FRK** | 10.1703 | 7.40E-05 | | **AMIGO1** | -12.0528 | 1.05E-05 |
| **SDCBP2-AS1** | 10.1637 | 5.22E-05 | | **NRIP3** | -12.0517 | 0.001078 |
| **TMEM51-AS1** | 1.02E+01 | 5.00E-05 | | **EDARADD** | -11.9542 | 0.002479 |
| **ZDHHC11** | 1.02E+01 | 4.34E-05 | | **MIR210HG** | -11.9286 | 0.000307 |
| **GSEC** | 10.1301 | 2.57E-05 | | **CBX2** | -11.9262 | 9.75E-05 |
| **CLCF1** | 1.01E+01 | 0.000484 | | **HOXD11** | -11.9154 | 0.004186 |
| **MYOM1** | 1.01E+01 | 0.002383 | | **SORD2P** | -11.9121 | 1.40E-06 |
| **B4GALT1-AS1** | 10.102 | 7.88E-05 | | **VLDLR** | -11.8932 | 0.000136 |
| **LOC101929066** | 10.0627 | 0.003347 | | **NAV2** | -11.8882 | 0.00022 |
| **STX1A** | 1.00E+01 | 0.000104 | | **CKMT1B** | -11.8666 | 0.000602 |
| **TFAP2A-AS1** | 1.00E+01 | 0.000192 | | **PAG1** | -11.8423 | 0.001896 |
| | | | | **PYGO1** | -11.828 | 0.000311 |
| | | | | **LOC154761** | -11.8055 | 0.002968 |
| | | | | **TMEM200A** | -11.8019 | 0.00223 |
| | | | | **LOC283299** | -11.7973 | 0.000151 |
| | | | | **TPRG1** | -11.7804 | 1.47E-05 |
| | | | | **PLA2G4A** | -11.7723 | 0.000168 |
| | | | | **GPX8** | -11.7434 | 2.10E-06 |
| | | | | **ST6GALNAC4** | -11.6894 | 8.58E-06 |
| | | | | **SCARA3** | -11.6649 | 5.55E-05 |
| | | | | **IFFO2** | -11.6251 | 0.000238 |
| | | | | **LDLR** | -11.5711 | 0.001532 |
| | | | | **CAMKK1** | -11.5593 | 0.000699 |
| | | | | **SLC39A2** | -11.5533 | 0.000149 |
| | | | | **BSPRY** | -11.5237 | 0.000123 |
| | | | | **CKMT1A** | -11.5083 | 0.000103 |
| | | | | **RGS14** | -11.4671 | 0.000183 |
| | | | | **PITPNM3** | -11.3599 | 0.000427 |
| | | | | **KREMEN1** | -11.3509 | 3.20E-05 |
| | | | | **ITPRIPL1** | -11.3229 | 0.002425 |
| | | | | **SPTLC3** | -11.1731 | 3.61E-05 |
| | | | | **FOLR3** | -11.1504 | 0.000584 |
| | | | | **INKA1** | -11.1333 | 0.001717 |
| | | | | **DNASE1L2** | -11.0445 | 0.000202 |
| | | | | **CYP2S1** | -11.0241 | 0.000241 |
| | | | | **RGS2** | -11.0048 | 0.002291 |
| | | | | **GJC3** | -11.0043 | 0.000966 |
| | | | | **TTC39A** | -11.0004 | 9.24E-05 |
| | | | | **CPT1C** | -10.9999 | 0.004209 |

| | | | | | | |
|---|---|---|---|---|---|---|
| | | | | **CBLC** | -10.9559 | 0.000131 |
| | | | | **IVL** | -10.9519 | 3.09E-05 |
| | | | | **ELOVL7** | -10.9104 | 0.000198 |
| | | | | **DHCR24** | -10.9058 | 1.05E-05 |
| | | | | **SOX7** | -10.8483 | 2.71E-05 |
| | | | | **PRRX2** | -10.7992 | 0.002522 |
| | | | | **RASGEF1A** | -10.7681 | 2.27E-05 |
| | | | | **HPGD** | -10.6973 | 0.002006 |
| | | | | **CEBPD** | -10.6916 | 0.000718 |
| | | | | **ZCCHC24** | -10.6879 | 0.000179 |
| | | | | **MGST1** | -10.6362 | 7.81E-05 |
| | | | | **FHDC1** | -10.5737 | 0.000228 |
| | | | | **CCNA1** | -10.528 | 0.000162 |
| | | | | **LYPD3** | -10.4425 | 4.51E-06 |
| | | | | **PDZD2** | -10.3565 | 0.000891 |
| | | | | **SCNN1G** | -10.3271 | 0.000121 |
| | | | | **HAS3** | -10.2834 | 0.000333 |
| | | | | **NKX1-2** | -10.2537 | 0.004397 |
| | | | | **ZFP42** | -10.2321 | 0.000783 |
| | | | | **EPGN** | -10.1995 | 0.000131 |
| | | | | **CARD18** | -10.1969 | 0.001112 |
| | | | | **STING1** | -10.1878 | 0.000902 |
| | | | | **DAPL1** | -10.1633 | 0.000302 |
| | | | | **PSG2** | -10.1367 | 0.000227 |
| | | | | **DMKN** | -10.1078 | 0.002369 |
| | | | | **WNT3** | -10.1057 | 5.94E-05 |
| | | | | **S100A2** | -10.0375 | 0.000668 |
| | | | | **TP63** | -10.0356 | 5.28E-06 |

**Supplementary Table 2 [**Related to **Figure 1H-J]:** Differentially expressed genes (DEGs) from SPT6-depleted vs. control samples from Li et al[1]. used to rank order normal (N) and Barrett's esophagus from human samples (BE) or wildtype (WT) and p63 knockout mouse samples in **Figure 1H-J**. Li et al[1] reported 472 upregulated genes (p-value < 0.01, >= 10-fold change), 528 downregulated genes (p-value < 0.01, >= 10-fold change).

| **Supplementary Table 3** [Related to **Figure 1K**] | | |
|---|---|---|
| **GSM ID** | GSE ID | Correlation Matrix ID |
| **GSM4634684** | GSE153129 | CTLi-1 |
| **GSM4634685** | GSE153129 | CTLi-2 |
| **GSM4634681** | GSE153129 | SPT6i-1 |
| **GSM4634682** | GSE153129 | SPT6i-2 |
| **GSM3415805** | GSE120795 | Colon-1 |
| **GSM3415821** | GSE120795 | Colon-2 |
| **GSM3415854** | GSE120795 | Brain-1 |
| **GSM3415888** | GSE120795 | Brain-2 |
| **GSM3415810** | GSE120795 | Ski mus-1 |
| **GSM3415864** | GSE120795 | Ski mus-2 |
| **GSM3701297** | GSE129153 | Adipocyte-1 |
| **GSM3701298** | GSE129153 | Adipocyte-2 |
| **GSM4483226** | GSE148818 | Trachea-1 |
| **GSM4483228** | GSE148818 | Trachea-2 |
| **GSM1423129** | GSE58963 | BE-1 |
| **GSM1423130** | GSE58963 | BE-2 |

**Supplementary Table 3 [**Related to **Figure 1K]**. Selection of GSM and GSE IDs used to generate the correlation matrix in **Figure 1K** including SPT6-depleted and control samples, colon, brain, skeletal muscle, adipocytes, trachea and Barrett's metaplasia.

| Supplementary Table 4 [Related to Figure 2A-C] | |
|---|---|
| **Metaplasia-specific and intestine-specific geneset** [PMID: 21703447] | |
| **Source:** https://doi.org/10.1016/j.cell.2011.05.026 | |
| **Metaplasia-Specific Genes:** | **Intestine-specific Genes:** |
| Cyp2f2 | Nr5a2 |
| Krt6a | Muc3 |
| Pax9 | Tinag |
| Adh7 | Afm |
| Upk2 | Hnf4a |
| Muc4 | Muc2 |
| Sox1 | Cdx1 |
| Gapbrp | Isx |
| Upk1a | Tff3 |
| Ceacam1 | Lgals2 |
| Cxcl17 | Cdx2 |
| Car4 | Alpi |
| Krt31 | Apob |
| Runx2 | Fabp2 |
| Krt7 | Lct |
| | Apoc3 |

**Supplementary Table 4 [**Related to **Figure 2A-C]**. Metaplasia-specific and intestine-specific genes used to perform GeneSet Enrichment Analysis (GSEA) on human Barrett's esophagus (BE) *vs*. small intestine tissue in **Figure 2B** and SPT6 KO *vs*. small intestine-derived organoids in **Figure 2C**.
